# Transcriptomic and Metabolomic Analysis of the Caudate and Putamen in ADHD

**DOI:** 10.64898/2026.11.06.764706

**Authors:** Rajvi Patel, Shuai Wang, Eshaan Deshmukh, Stephen Fajen, Jamal Williams

## Abstract

Attention-deficit/hyperactivity disorder (ADHD) is a highly heritable, heterogeneous neurodevelopmental disorder that typically manifests in childhood; however, transcriptomic studies have inadequately accounted for age of onset and subtype, with cohorts largely comprising of adults. To address this gap, we established a stringently phenotyped ADHD cohort (age ≤19 years; clinically diagnosed combined and inattentive-type ADHD) and integrated postmortem caudate and putamen bulk RNA-seq, LC-MS metabolomics, MEGENA gene co-expression analysis, and cross-study comparison. We identified 279 up- and 35 downregulated DEGs in the caudate and 110 up- and 89 downregulated DEGs in the putamen. The combined caudate-putamen metabolomic analysis revealed dopamine dysregulation (1.68-fold reduction) and dysregulation of four additional metabolites. Enrichment analysis revealed shared molecular signatures involving immune responses, ATP metabolism and Ca²⁺-dependent signaling, with region-specific differences in the direction of alteration. Neuronal homeostasis, and protein stress showed suppression in the putamen, along with activation of neurotransmitter reuptake, whereas CSF circulation and ECM organization were activated in the caudate, with suppression in CNS myelination, and signs of ventricular perturbations. MEGENA identified 26 ADHD-associated caudate modules and 33 putamen modules. These modules captured canonical synaptic and neuronal-circuitry pathways alongside cellular differentiation, metabolism, and cell-survival processes. FGD1 emerged as a candidate caudate hub gene among the neuronal modules. Cross-study comparison analysis suggested that molecular alterations in ADHD may be age-dependent. Additionally, assessing levels of 76 potential risk genes’ identified by a large ADHD GWAS study one ARTN exhibited dysregulated levels in putamen and none in caudate. Together, these findings reveal brain-region-specific molecular dysregulation in ADHD across pathways extending beyond canonical neurotransmission to immune, differentiation, and energy-metabolic processes, along with novel evidence of perturbations in the understudied lateral ventricles.

## Introduction

Attention-deficit/hyperactivity disorder (ADHD) is a common neurodevelopmental disorder (NDD) that is largely characterized by symptoms of inattention, hyperactivity or impulsivity. ADHD has a high heritability of ∼80%^1^ and affects 11.4%^1, 2^ of school-aged children in the United States and 7-8% internationally^3^, with the prevalence rates increasing every year^4^. While earlier foundational research focused on investigating structural and functional alterations among brain regions^5, 6, 7, 8^, the recent advances in genomics over the past few decades have shifted to focus towards identifying the underlying genetic factors of the disorder through genome-wise association studies^9, 10^. These studies have also helped in discovering imputed genes associated with ADHD^11, 12^. Transcriptomic studies conducted with peripheral blood tissue identifying alterations in networks related to nervous system development, lipid metabolism and galactose transcriptome^13, 14^. While all these studies provide highly valuable and informative knowledge in understanding several aspects of the disorder, very few studies have conducted expression-level studies identifying the levels of genes and metabolites and the studies that have been conducted are either using animal models or using human blood and urine samples giving indirect means of understanding the underlying cellular and molecular mechanisms altered in the human brain in ADHD^11, 13, 14, 15^. So, the biggest gaps remain (a) in the link between GWAS-identified SNPs and imputed genes to identifying genes with altered expression levels in human tissues, (b) from indirect sources of expression to finding the genes and metabolites with dysregulated expression levels in the brain, and (c) identifying the consequential alteration in biological pathways that underlie symptoms and pathophysiology of ADHD. Our study aims to address these gaps by utilizing post-mortem brain tissue, identifying the dysregulated genes and metabolites, and elucidating their consequential cellular and molecular functions in 2 specific brain regions, namely the caudate and putamen.

Another big gap comes from the heterogeneity of this disorder at several fronts, including age of onset, presentation of symptoms across and within each subtype, different brain regions and circuits being affected differently between each subtype, causal factors^1^, and presented comorbidities^16^. Despite these, the research so far has been conducted at a very generalized scale, which helps in increasing the statistical power of the study but results in findings that are biased toward the most represented group in the study. This is a potential reason underlying the variability seen between the findings from different studies^17^ and leads to loss in our ability to find accurate mechanisms. Thus, our study design focuses on reducing the heterogeneity by implementing a highly selective phenotyping criteria focusing on childhood-adolescence age group (<= 19 years), clinically diagnosed (no self-reported) inattentive and combined type ADHD individuals, being inclusive of both genders and excluding individuals with schizophrenia as a comorbidity.

Our focus on this age-group is for several reasons. Firstly, ADHD is a NDD, so the physiological and molecular alterations likely begin to manifest as early as during infancy or prenatal stages, much before clinical diagnosable symptoms onset. Secondly, the peak age of diagnosis is 9.5 years, with a median of 12 years and 73.0% of individuals receiving the diagnosis by age of 18^1, 18^. For a disorder with such an early onset, it is crucial to investigate the pathophysiological mechanisms underlying this developmental period. Thirdly, studies have reported an overall developmental delay in the brain, reporting an overall 3% reduction in volume. This reduction in brain volume has been reported individually^19, 20^ for several brain regions, such as the corpus callosum^21, 22, 23^, prefrontal cortex^24, 25^, basal ganglia^25, 26, 27^, cerebellum^28, 29^ and straitum^20, 30, 31^. These differences are more prominent in children, but do not persist consistently with age depending on brain region^19, 20^.

Neuroimaging, neurochemical, genetics, and neuropsychological studies have implicated the fronto-striatal network, comprising of lateral prefrontal cortex, dorsal anterior cingulate cortex, putamen and caudate, in ADHD pathophysiology^32, 33, 34^. Among structural findings, reduced caudate volume has been one of the most consistently reported alterations^20, 30, 31^ while reduced putamen volume has also been frequently observed^20, 30, 35^. Several studies have further reported altered activation of the caudate and putamen, as well as their altered connectivity with other brain regions in ADHD^34, 36, 37^. Caudate and putamen together form the dorsal striatum, which plays a critical role in reward processing, action selection, and decision-making^38^, functions that are commonly affected in ADHD. However, the dorsal striatum is both molecularly and functionally heterogeneous. The caudate is preferentially integrated with associative and prefrontal networks supporting flexible, goal-directed and cognitively controlled behavior, whereas the putamen is strongly connected with sensorimotor networks supporting movement, motor learning and habitual action. These functional differences stem from differences in their neuronal connectivity, circuitry they are part of, cell-type composition, and their cellular and molecular processes^39, 40, 41, 42^. Accordingly, transcriptomic and metabolomics alterations in caudate and putamen may also differ in ADHD, despite both regions belonging to the same striatal system.

We tested three major hypotheses: (1) the caudate and putamen harbor key transcriptomic and metabolic alterations associated with ADHD; (2) despite belonging to the same striatal system, the two regions exhibit distinct molecular alteration profiles; and (3) these alterations vary with the developmental age at which ADHD is presented. Here, we conducted bulk RNA sequencing on post-mortem brain tissue from the caudate and putamen, identified the up- and down-regulated genes in and within these brain regions, and conducted a metabolomic study identifying the metabolites with altered levels. We identified 5 metabolites, including well-identified dopamine levels, to be altered in both brain regions. In line with our hypotheses, we found that out of 757 highly expressed, significantly differentially expressed genes (p-value <=0.05 and fold-change of >= |2|), only 20 genes overlapped between the two regions, accounting for only 2.64% mRNA-level commonality. Several novel pathways were identified to be dysregulated at cellular and molecular levels, including but not limited to immune and inflammatory response, ATP metabolism, intracellular vesicle transport, neurotransmitter availability, and strong signs of developmental delay in one or both brain regions. Of these, the first two exhibited opposite directional dysregulation between caudate and putamen, in line with our first hypothesis.

We also conducted cross-study comparative analysis with 2 major studies. We compared our age-specific ADHD associated DEGs in caudate with the only other existing transcriptomic study in the field and its age-independent ADHD associated caudate DEGs,^43^ where we observed an inverse correlation between the overlapping DEGS from the two studies. Lastly, we looked for the expression of 76 potential risk genes associated with the 27 genetic loci identified by one of the largest GWAS studies by Demontis et al.^12^ and found that only one gene ARTN was found to have altered expression, in putamen and none showed dysregulation in caudate. This demonstrates that disorder-associated SNPs are not differentially expressed in all tissue types, emphasizing the need for more transcriptomic and molecular studies to advance the field of ADHD research. While our analyses reveal meaningful findings, the limited sample size (ADHD = 6 and Control = 3 individuals) owing to our rigorous phenotyping criteria, especially at early developmental age range, comes with the caveat of potentially reduced statistical power and sensitivity of detecting alterations.

## Results

### Cohort design and characterization as key foundation of the project

A major knowledge gap exists connecting the findings from GWAS to gene expression data from human tissue, and importantly an insufficiency in later. To address the latter we investigate the transcriptomic profiles of 2 striatal brain regions, the caudate and putamen. To minimize phenotypic heterogeneity, we first established stringent cohort-selection criteria. These criteria restricted the ADHD and control cohorts to individuals aged 10–19 years, consistent with the World Health Organization’s (WHO) age range of adolescence. The ADHD cohort was further restricted to individuals with a clinically confirmed diagnosis of combined-subtype ADHD, excluding self-reported cases and individuals with schizophrenia as a comorbidity. Application of these stringent criteria substantially reduced the number of eligible biobank samples with final cohorts comprising of 6 ADHD individuals and 3 control individuals. Both male and female individuals were included to ensure representation of both sexes. Additionally, to reduce inter-cohort variability between the caudate and putamen analyses, the two brain-region cohorts were composed largely of the same individuals, with only one unique individual in each cohort (93052 in caudate’s and 93027 in putamen’s control cohorts). These details of individuals that comprised these cohorts can be found in Supplementary Table 1.

RNA transcripts from samples of all four cohorts were sequenced, followed by processing of the raw reads and post-quality filtering with fastp, where caudate samples retained an average of 41.3 ± 7.0 million reads per samples (min=33.0 and max+=53.2 million reads per sample) and putamen samples retained 47.3 ± 8.4 million reads per sample (min=37.9 and max=61.5 million reads per sample). The principal component analysis (PCA) of caudate’s normalized counts showed PC1 and PC2 accounted for 47% and 16% of variance respectively and PC1–PC4 together captured ∼80% of the total variability in the expression dataset (Supplementary Fig. 1A). PC1and PC2 accounted for 37% and 17% of variance respectively in putamen’s dataset with PC1-PC5 capturing ∼80% of variability (Supplementary Fig. 1B).

### Dorsal striatum’s altered transcriptome in ADHD

Significantly differentially expressed genes (DEGs) were identified from the bulk RNA-sequencing data as the genes that have gene expression significantly different with p-value <=0.05 (Fig. 1A & B, horizontal dotted line) in ADHD cohort as compared to their expression in the control cohort. From these, DEGs with gene expression higher or lower than a fold change of 2 (Fig. 1A & B, vertical dotted line) in ADHD cohort as compared to control cohort, were described in this paper as highly expressed significantly differentially expressed genes (HES-DEGs). 432 genes were identified as significant DEGs of which 346 were up-regulated and 86 were down-regulated and 279 were up-regulated HES-DEGs and 35 were down-regulated HES-DEGs (Supplementary Table 2, Fig 1A). Putamen identified a total of 347 significant DEGs almost equally distributed between up- and down-regulated directions, 174 being up-regulated and 173 being down-regulated, and of these 110 are up-regulated HES-DEGs and 89 are down-regulated HES-DEGs (Supplementary Table 3, Fig 1B).

**Figure 1:**
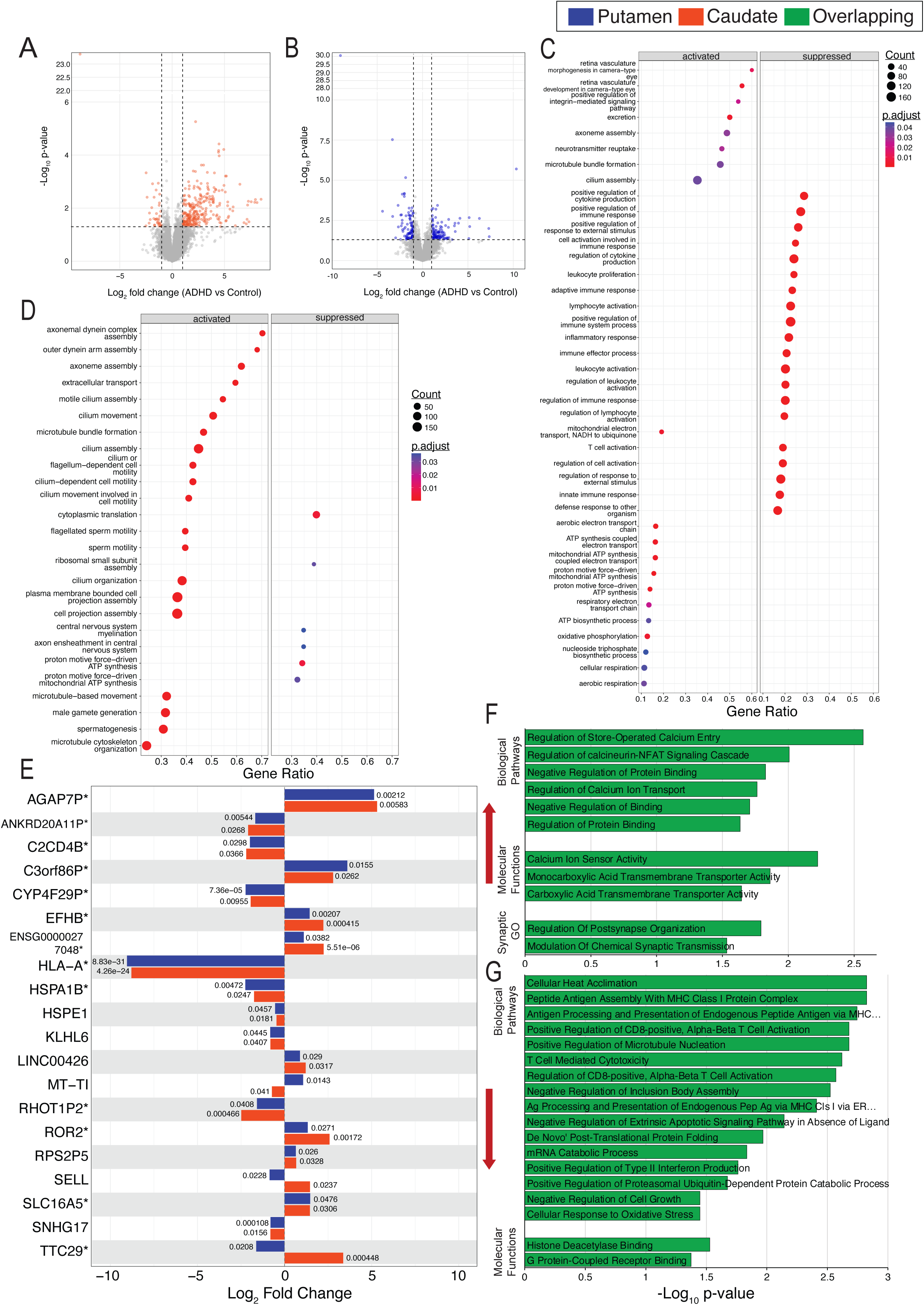
ADHD-associated differentially expressed genes in Caudate and Putamen and their potential downstream consequences: (A,B) Volcano plots of Bulk RNA-sequencing identifying the differentially expressed genes (DEGs) in ADHD group as comapred to Control group individuals in Caudate (A) and Putamen (B) tissues (n: ADHD=6, Control=3). The colored dots in the plots (organge in Caudate and blue in Putamen) highlight DEGs with p-value =<0.05 and fold change >= 2 (marked with dotted horizontal and vertical threshold lines respectively). This paper defines DEGs above these thresholds as Highly Expressed Significant DEGs (HES-DEGs). (C, D) GSEA analysis of the DEGs from Putamen (C) and Caudate (D) identifying the Biological Processes that these DEGs would cumulatively affect along with the direction of their effect i.e. in up-(activated) or down-(suppressed) regulated manners. (E) Significant DEGs expressed in both Caudate and Putamen with their expression (Log2FoldChange on x-axis) in each of these brain regions individually and their p-values mentioned besides their corresponding bars. From these overlapping significant DEGs, the genes with a higher differential expression i.e., FoldChange >= 2 (i.e., HES-DEGs) are marked with *. (F) Biological Processes, Molecular Fucntions and Synaptic components (as per synaptic gene onltology (GO)) associated with the up-regulated overlapping HES-DEGs as identified using EnrichR. (G) Biological Processes and Molecular Fucntions associated with the down-regulated overlapping HES-DEGs as identified using EnrichR.

#### Caudate’s transcriptomic landscape

Together, the significant DEGs in caudate revealed to predominantly govern activation of cytoskeletal cilia assembly and movement by biological processes (BPs) such as axonemal and outer dynein complex assembly, microtubule bundle formation, cilium movement, flagellum dependent cell motility, and others that were identified to be activated upon conducting Gene set enrichment analysis^44, 45^ (GSEA) (Fig. 1D, Supplementary Table 4). All top 20 activated BPs from GSEA analysis pertained to ciliary assembly and motility (Fig. 1D), with some BPs being associated with ∼150 DEGs (Fig. 1D), and 92 DEGs were associated with the top 5 BPs (Supplementary Fig. 2C, Supplementary Table 4). In addition, BPs such as cerebrospinal fluid (CSF) circulation, response to retinoic acid, pattern specification, as well as immune system and immune cell regulation, activation, and proliferation were also found to be activated (Supplementary Table 3). On the other hand, these DEGs are associated with suppression of cytoplasmic translation, CNS myelination and proton motive force−driven ATP synthesis (Fig 1 D). Analyzing the molecular functions (MFs) associated with these significant DEGs in caudate reveal involvement with activation of immune and signaling receptor activities, growth factor binding, in addition to, microtubule motor activity, dynein’s chains bindings and extracellular matrix (ECM) structural constituent, consistent with the activation of several cilia-related molecular functions. Along with suppression of structural constituent of ribosome, NADH dehydrogenase (ubiquinone) and oxidoreduction-driven transmembrane transporter activities (Supplementary Fig. 2A, Supplementary Table 5).

#### Putamen’s transcriptomic landscape

While putamen’s GSEA analysis revealed significant (padj. <= 0.05) suppression in immune response and immune cells by suppression of BPs including, as but not limited to, positive regulation of cytokine production, leukocyte proliferation, leukocyte, lymphocyte activation and T-cell activations, immune effector process and defense response to other organism, with >=120 DEGs associated with certain individual BPs such as response to external stimulus (Fig 1D, Supplementary Table 4). All top 20 suppressed BPs were found to be involved in immune response and activation (Fig. 1E), with 204 DEGs being associated with the top 5 BPs (Supplementary Fig 2D, Supplementary Table 6). Additionally, other BPs including learning, gliogenesis, neuron remodeling, negative regulation of neurogenesis and positive regulation of cell migration, proteolysis and hydrolase activity and more were also identified to be suppressed (Supplementary Table 6). These DEGs were also found to be related to activation of a broader variety of BPs related to morphogenesis, positive regulation of integrin-mediated signaling pathway, neurotransmitter reuptake, mitochondrial electron transport (NADH to ubiquinone), in addition to 2 BPs related to cilium assembly and several BPs related to (mitochondrial) ATP synthesis i.e., mitochondrial/ cellular respiration. These DEGs also imply suppression of mechanisms that regulate and respond to cellular protein stress by suppression of MFs such as heat shock protein binding and protein folding chaperone, along with suppression of cysteine type peptidase activity, transmembrane signaling receptor activity and molecular transducer activity. NADH dehydrogenase activity, actin binding, electron transfer activity, oxidoreductase activity acting on NAD(P)H, quinone and similar compound and Wnt receptor activity were potentially activated MFs (Supplementary Fig. 2B, Supplementary Table 7).

#### Similarities and differences between caudate and putamen’s transcriptome

Despite the several differences in the BPs associated with caudate and putamen’s DEGs, two notable region-specific patterns emerged. Both brain regions’ reported BPs associated with immune response/ activation, however in opposite direction, with suppression in putamen and activation in caudate. Similarly, BPs related to ATP synthesis were also seen to be conversely correlated in both brain regions with suppression in caudate and activation in putamen.

Among the 757 significant DEGs identified across the two brain regions, only 20 were shared between the caudate and putamen, representing 2.64% overlap in differential gene expression (Fig. 1E, Supplementary Table 8) and 3 DEGs have expression in opposite direction (SELL, TTC29 and MT-TI). Of these 20, 13 genes have a fold change >= |2| (Supplementary Table 8), i.e., overlapping HES-DEGs, with 6 genes being up-regulated, 6 genes being down-regulated, and 1 gene (TTC29) having expression in the opposite direction in the two brain regions (Fig. 1E, marked by *, Supplementary Table 8). These shared up- and down-regulated genes were analyzed for their functional enrichment on the EnrichR^46, 47^ platform using Biological Process, Molecular Function and Cellular Component Gene Ontologies^48^ as well as Synaptic Gene Ontology^49^ to identify the shared dysregulation between the caudate and putamen brain regions. Analysis of the shared up-regulated, identified these DEGs to be are involved in calcium ion movement and its dependent pathways by regulating store-operated calcium entry, calcium ion transport, calcineurin-NFAT signaling cascade along with negative regulation of protein binding biological processes and molecular functions involved in Calcium ion sensor activity and cellular metabolism via monocarboxylic acid transmembrane activity (Fig. 1F, Supplementary Table 9). Their synaptic GO analysis identified enrichment in regulation of post-synapse organization and modulation of chemical synaptic transmission (Fig. 1F) and Cellular component enrichment analysis identified clathrin-coated endocytic vesicle–related compartments, along with motile cilia/flagellar components (Supplementary Table 9). Whereas the shared down-regulated DEGs showed enrichment in several BPs of immune response especially via MHC Class I complex, T-cell, CD4 and 8 regulation, alongside protein stress and quality-control pathways, cellular responses to oxidative stress, regulation of cell survival/ growth via negative regulation of apoptotic signaling pathways, mRNA catabolism, and microtubule organization (Fig. 1G, Supplementary Table 9). These DEGs are also involved in molecular functions) involved in binding of histone deacetylase and G-protein coupled receptor (Fig. 1G, Supplementary Table 9). Cellular Component enrichment of these overlapping down-regulated DEGs highlighted compartments involved in ER–Golgi trafficking and vesicular transport, including the Golgi medial cisterna, ER-to-Golgi transport vesicle membrane, coated vesicle membrane, and transport vesicle membrane, together with enrichment of the lumenal side of the ER membrane. Immune- and degradation-related compartments, including MHC protein complexes, phagocytic vesicle membrane, and aggresome, were also enriched (Supplementary Table 9).

### Several genes share co-expression patterns with a strong correlation to ADHD, in caudate and putamen brain tissue

To identify coordinated gene-expression patterns beyond individual DEGs, we applied Multiscale Embedded Gene Co-expression Network Analysis (MEGENA)^50^ to the top 2,500 ranked DEGs (ranked by (-log_10_p-value) × absolute(log_2_fold-change)) from the caudate and putamen individually. MEGENA groups genes with correlated expression profiles into co-expression modules, allowing both significantly and non-significantly dysregulated genes to contribute to network-level patterns. Because genes function within coordinated regulatory networks rather than in isolation, dysregulation of individual genes can influence the expression and function of other genes in the network. Including both significant and non-significant genes therefore enables identification of broader co-expression patterns and biological processes that may not be captured by DEG-focused analyses alone. MEGENA organizes these modules hierarchically, with broad parent modules progressively branching into smaller child modules containing genes with increasingly similar expression patterns and concluding in leaf modules at the end of these branches. Upon identification of modules, they were then tested for association with ADHD. 177 co-expression modules were identified in caudate of which 26 modules were significantly associated with ADHD (t-test, p ≤ 0.05), with 11 parent and 15 leaf modules (Supplementary Table 11; Fig. 2A, zoomed in hierarchal plots in Supplementary Fig. 4A). 188 modules were identified in the putamen of which 33 modules were significantly associated with ADHD (t-test, p ≤ 0.05), with 10 parent and 23 leaf modules (Supplementary Table 12; Fig. 2C, zoomed in hierarchal plots in Supplementary Fig. 5A). Module eigengene scores (ME) were also calculated for each module. This value represents the summarized overall expression pattern of all genes within a module across samples. The ME value showed distinct expression patterns between control and ADHD samples for all modules in both brain regions (Supplementary Fig. 3A & B). All caudate modules had a lower ME in ADHD samples than in controls (Supplementary Fig. 3A, Supplementary Table 10), indicating coordinated reduced expression pattern of gene of modules in ADHD cases compared with controls. In addition, all modules of caudate showed a significant negative correlation (calculated by Pearson correlation) with ADHD phenotype (Supplementary Table 11). On the other hand, 10 putamen modules showed higher ME in ADHD as compared to control, indicating a relatively higher coordinated gene expression pattern in ADHD as well as a significant positive correlation to ADHD phenotype, and 22 modules showed a relatively reduced coordinated gene expression pattern (by a reduced ME score in ADHD samples as compared to controls) and a significant negative correlation to the ADHD phenotype (ME score: Supplementary Fig. 3B, Supplementary Table 10, r: Supplementary Table 14).

**Figure 2:**
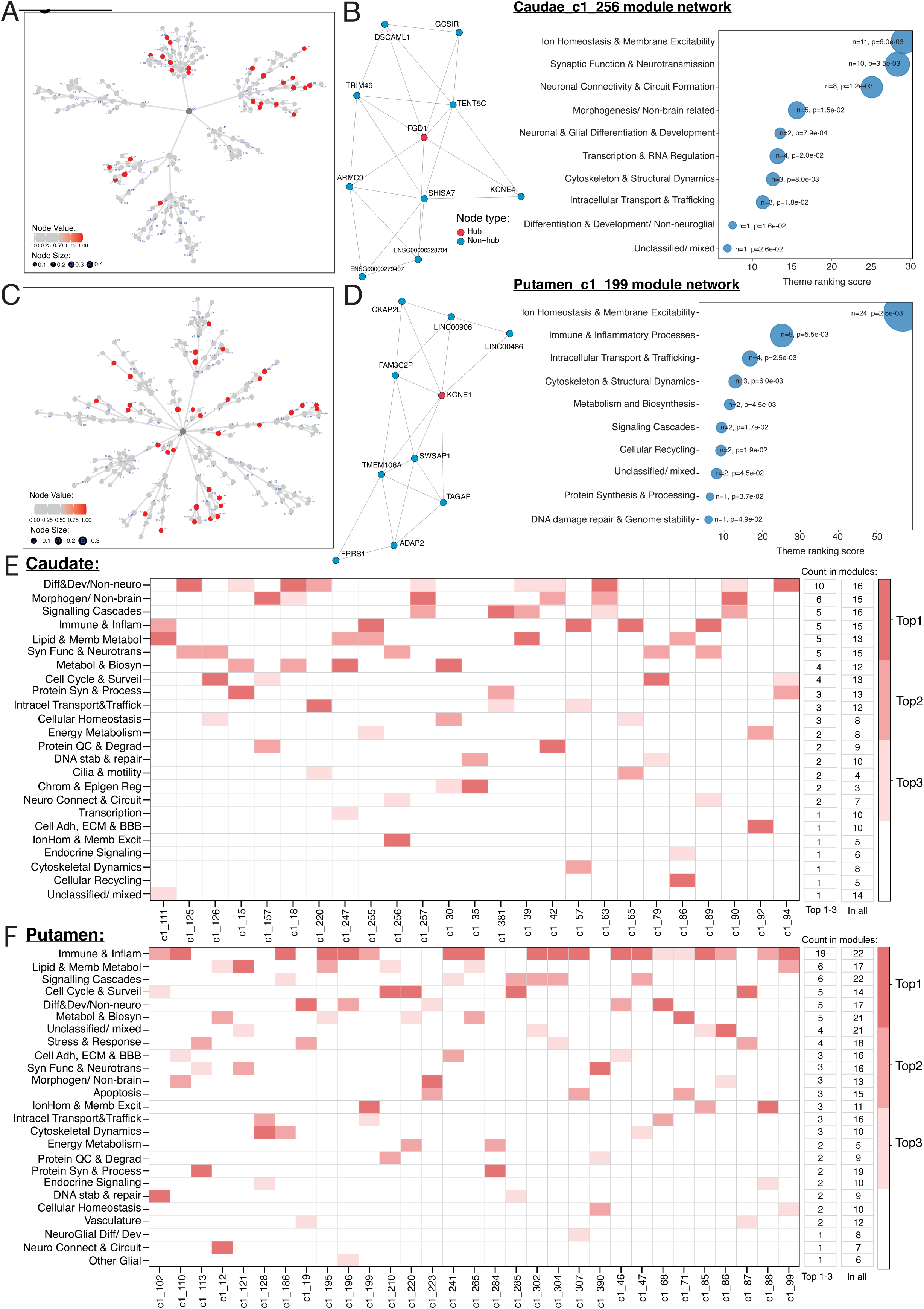
Gene Co-expression Network Analysis of Caudate and Putamen DEGs: (A, C) Hierarchical plots displaying the entire network of modules constructed through Multiscale Embedded Gene Co-expression Network Analysis (MEGENA) analysis of top 2500 ranked DEGs of caudate (A) and putamen (C). Each module consists of genes grouped together based on their co-expression patterns in Control and ADHD groups combined. Red nodes represent modules that are significantly associated with ADHD phenotype (p-value <= 0.05) and size of node represents the number of genes in the module i.e. module size. (B & D) Module c1_256 and c1_199 from caudate and putamen repsectively, represent zoomed in version of modules displaying their network plots with the genes constituting the module, the connections between those genes and the Hub gene (highlighted in red) that are the highly connected genes within the module; followed by the dot plot exhiting the biological themes (functions) associated to the module’s genesets. Each biologocal theme classifies the very specific and precise Gene Ontology (GO) terms of Biologiocal Processes (BPs) from EnrichR results, into broader and interpretable terms. The themes are ranked by a ranking metric of: Rank = 3(-log10p-value) + 2n + combined score, where n is the number of GO terms in the theme, -log10p-value of the top BP in the theme and its combined score both as calculated by EnrichR. (E, F) Heatmaps representing all the modules of Caudate and Puatemen respectively and theTop 3 biological themes assocaited with each module’s genesets. The biological themes are represented in acronyms with the guide to their full names mentioned in Appendix/ Table #. The columns on the right of the heatmap mentions, for every theme: the number of modules for which that theme is a top (1^st^ to 3^rd^) ranking theme (first column) and the number of modules that have that biological theme (as its associaed function) irrespective of the rank (second column). [GO terms are widely used structured, standardized representation of biological knowledge/ terms (cite GO’s website) and EnrichR identifies Biological Processes in the form of GO terms]

### MEGENA provides more evidence for biological dysregulation identified from DEGs’ enrichment analyses by revealing consistent functional consequences from ADHD-associated gene modules

To extend the analysis beyond module identification, these significantly correlated modules were then evaluated for their functional consequences, using EnrichR’s^46, 47, 51^ Gene Ontology Biological Process^48^ (GO-BP) enrichment analysis to identify biological processes associated with the genes within each module. As GO-BP enrichment analysis often yields numerous highly specific and overlapping GO terms, we further summarized the GO terms into 30 manually curated functional themes, using specific curated definitions for summarization (Supplementary Table 16). This additional classification helped reduce redundancy from the multiple GO terms, summarized them into broader, interpretable biological functions, and enabled functional annotation of each module along with identification of the dominant/ representative biological functions it governs.

The gene(s) with the highest connectivity to the rest of the genes within that module is identified as the module’s hub gene(s), for instance FGD1 and KCNE1 are the hub genes of modules c1_256 and c1_199 in the caudate and putamen, respectively (Fig. 2B, D). All modules reported associations with several functional themes (Supplementary Table 17, 14). So the functional themes were ranked to identify the dominant functions of each module. The rank was calculated as: Rank = 3(-log10p-value) + 2n + combined score (where n is the number of GO terms in the theme, -log10p-value and combined score are of the top BP in the theme; all these as were reported by EnrichR for the module’s gene set). The top 10 functions were plotted in the dot plots, with dot size proportional to the number of GO terms present for that theme (Fig. 2B, D). The ranking also helped identify specific modules related to specific functions such as ‘cell cycle & surveillance’, ‘protein QC, stress and degradation’, as well as neuronally centered functions such as ‘synaptic function and neurotransmission’, ‘ion homeostasis and membrane excitability’, ‘Neuronal & Glial Differentiation & Development’, and many more (Fig. 2E, F).

Several modules’ functions were found to be consistent with DEG’s GSEA analysis. In line with putamen’s DEGs’ GSEA analysis that identified suppression of immune response, putamen’s MEGENA analysis also reported 22 (out of 33) modules with ‘immune & inflammatory processes’ as their function and 19 modules reporting it as one of their top 3 functions (Fig. 2F, Supplementary Fig. 5B). c1_186 and c1_99 reported this as its top function with a rank of >120 and >50 respectively (Supplementary Fig. 9C, E).

The suppression of ATP synthesis in caudate and its activation in putamen through GSEA-BP analysis, MEGENA revealed ‘Energy metabolism’ to be a top 1-3 function of 2 modules in caudate as well as in putamen with negative correlation to ADHD. Also, in line with caudate’s suppression of cytoplasmic translation, 3 modules were associated with ‘Protein Synthesis & Processing’ as the top 1-3 function. Table 1 summarizes a list of functions identified through both brain regions’ DEGs’ enrichment analyses (GSEA BP and MF analysis and EnrichR’s analyses for the shared DEGs’) and their corresponding MEGENA functional themes. For each theme, the table reports the number of modules associated with that functional theme and the number of modules for which that functional theme is ranked among the top three functions (Fig. 2E-F; Supplementary Fig. 4B, 5B).

**Table 1:** Correlation between functional consequences identified from DEGs and from Gene co-expression modules:

| Brain Regions: | Direction of dysregualtion: | GSEA/ EnrichR- BP/ MF/ SynGO: | Correlating MEGENA's functional theme: | # of modules with this fucntion amongst it top 3 fucntions: | # of modules with this as its fucntion: |
|---|---|---|---|---|---|
| Common amongst both regions' individual GSEA analyses' | Cau: up, Put: down, shared DEGs: down | Immune reponse and activation | Immune & Inflammatory Processes | Cau: 5; Put: 19 | Cau: 15; Put: 22 |
|  | Cau: down, Put: up | ATP synthesis | Energy Metabolism | Cau: 2; Put: 2 | Cau: 8; Put: 5 |
| Shared DEGs' (between Cau and Put) EnrichR analysis | up | calcium ion trnasport | Intracellular Transport & Trafficking, Synaptic Function & Neurotransmission, | Cau: 3, 5; Put: 3, 3 | Cau: 12, 15; Put: 16, 16 |
|  |  | (calcium dependent) calcineurin-NFAT signalling pathway | Signaling Cascades, Immune & Inflammatory Processes | Cau: 5, 5; Put: 6, 19 | Cau: 16, 15; Put: 22, 22 |
|  |  | post-synapse organization | Neuronal Connectivity & Circuit Formation | Cau: 2; Put: 1 | Cau: 7; Put: 7 |
|  |  | modulation of chemical synaptic transmission | Synaptic Function & Neurotransmission | Cau: 5; Put: 3 | Cau: 15; Put: 16 |
|  |  | cellular metabolism | Metabolism and Biosynthesis | Cau: 4; Put: 5 | Cau: 12; Put: 21 |
|  | down | protein stress and quality control | Preotein QC, Stress & Degradation | Cau: 2; Put: 2 | Cau: 9; Put: 9 |
|  |  | cellular stress and growth | Stress & Stimulus Response, Apoptosis | Cau: -,- ; Put:-, 3 | Cau: 12; Put: 18 |
|  |  | mRNA catabolism | Transcription & RNA Regulation | Cau: 1; Put: - | Cau: 8; Put: 12 |
| Caudate's GSEA analysis | up | ciliary assemby and motility | Cilia & Ciliary motility | 2 | 4 |
|  |  | CSF circulation | Cell Adhesion, ECM & Barrier Integrity | 1 | 10 |
|  |  | signaling receptor activities | Signaling Cascades | 5 | 16 |
|  |  | ECM structural constituent | Cell Adhesion, ECM & Barrier Integrity | 1 | 10 |
|  | down | cytoplasmic translation | Protein Synthesis & Processing | 3 | 13 |
|  |  | CNS myelin | Other Glial Processes and Functions | - | 1 |
|  |  | NADH dehydrogenase | Energy Metabolism | 2 | 8 |
|  |  | oxidoreduction-driven transmembrane transporter activities |  |  |  |
| Putamen's GSEA analysis | up | neurotransmitter reuptake | Other Glial Processes and Functions | 1 | 6 |
|  |  | cellular respiration | Energy Metabolism | 2 | 5 |
|  |  | wnt receptor activtiy | Signaling Cascades | 6 | 22 |
|  |  | integrin-mediated signaling pathway | Cell Adhesion, ECM & Barrier Integrity | 3 | 16 |
|  |  | morphogenesis | Morphogenesis/ Non-brain related | 3 | 13 |
|  | down | proteolysis | Preotein QC, Stress & Degradation | 2 | 9 |
|  |  | cellular preotein stress and response |  |  |  |
|  |  | hydrolase activity |  |  |  |
|  |  | cell migration | Neuronal Connectivity & Circuit Formation | 1 | 7 |
|  |  | signaling receptor activity | Signaling Cascades | 6 | 22 |

Cau= Caudate; Put= Putamen

### MEGENA revealed several overshadowed neuronal functional consequences from ADHD-associated co-expression modules in the Caudate, along with identifying a functionally neuronally enriched module

MEGENA also identified multiple additional functions associated with ADHD, along with their corresponding modules and gene sets, that may get overlooked in DEG-focused analyses because they are either overshadowed by more dominant DEG-enriched pathways or depend on coordinated contributions from non-significantly dysregulated genes that co-express with significant DEGs. For instance, in addition to identifying 4 of modules functionally associated with ‘Cilia and Motility’ in caudate (Supplementary Fig. 4B), with 2 modules having it as their top 3 ranked function (Fig. 2E), consistent with the caudate’s predominantly identified BP from DEGs’ GSEA analysis (Fig. 1D), MEGENA analysis also reported ‘Synaptic Function & Neurotransmission’ as functions of 15 modules, ‘Neuronal connectivity and Circuit Formation’ of 7 modules and 6 modules were identified to be associate with ‘Neuronal & glial Differentiation & Development’ (Supplementary Fig 4). Functionally, 6 modules were identified to be associate with ‘Neuronal & glial Differentiation & Development’, 5 with ‘Ion Homeostasis and Membrane Excitability’, 5 with ‘Vasculature and Angiogenesis’, 10 ‘Cellular Adhesion, ECM and BBB integrity’, 1 with ‘Brain Morphogenesis’ and 1 with ‘Other Glial Processes and Functions’ (Supplementary Fig 4), along with several modules reporting these in their top3 functions (Fig 2). However, the top 3 functional themes captured the largest proportion of GO terms associated with the module and showed the strongest statistical significance (lowest p-values),for most modules, as can be observed from the ranked functional profile of module c1_256’s (Fig. 2B dot plot). Therefore, the top 3 ranked functional themes were considered to define the module’s function as the representative (or dominant) function of the module and for subsequent downstream analyses.

5 modules had ‘Synaptic Function & Neurotransmission’ as one of their dominant functions, which annotates biological processes (BPs) related to neurotransmitter availability, from their production and transport to their exocytosis at synapse aided by calcium ions. 2 modules had ‘Neuronal connectivity and Circuit Formation’ as one of their dominant functions, and it annotates BPs involved in organization and development of neural circuitry including axongenesis, neuron projection guidance, guidance cues and related processes. In addition, ‘Cell Adhesion, ECM & Barrier Integrity’ that annotates BPs related to cell-cell and cell-matrix interaction, ECM organization and maintenance of the blood brain barrier (BBB) and ‘Ion Homeostasis & Membrane Excitability’ that annotates BPs related to homeostasis, including transport, channel activities of Na+, K+, or Cl-(except Ca+2) ions, and their consequential membrane excitability states, are dominant functions of 1 module each. (Supplementary Table 16 shows how each functional theme was defined, and Fig. 2E shows the number of modules for which each functional theme is dominant.)

Owing to MEGENA’s hierarchical characterization of modules, the terminal leaf modules attain functional specification as compared to their parent modules. The family of caudate’s c1_89 (parent), c1_255 (leaf) and c1_256 (leaf) modules, where c1_89, the parent module, has 21 biological themes, whereas the leaf modules c1_255 and c1_256 have 18 and 12 biological themes respectively associated with their functional consequence (Supplementary Fig. 6E). Most of their biological themes overlap with each other, 7 overlapping between all three of them, including ‘Synaptic Function & Neurotransmission’, ‘Neuronal Connectivity & Circuit Formation’, ‘Signaling Cascades’, ‘Cytoskeleton & Structural Dynamics’ and 3 other (Supplementary Fig. 6F). The functional specification is reflected in the co-expression pattern specification also, with leaf modules consisting of fewer genes than their parent module (Fig. 2B and Supplementary Fig. 6B, D: network plots, Supplementary Table 11: module size; Supplementary Table 14: gene list).

Module c1_89 is associated with 2 neuronal functional themes as its dominant functions, related to neurotransmitter availability, i.e., ‘Synaptic Function & Neurotransmission’, and neuronal circuit development, i.e., ‘Neuronal Connectivity & Circuit Formation’ (Supplementary Table 16, Supplementary Table 17). Module c1_126, which is also a parent module, is associated with neuronal function related to neurotransmitter availability as its dominant function, along with ‘Cell Cycle & Surveillance’ and ‘Cellular Homeostasis’ (Supplementary Fig. 7A, Supplementary Table 12). Module c1_256 was particularly notable for its concentrated association with neuronal functions, with all three top-ranked dominant functional themes corresponding to neuronal processes. It consisted of 10 genes with FGD1 being the central regulatory gene i.e. hub gene of the module (Fig. 2B, Supplementary Table 14).

In addition to modules governing several neuronal processes, MEGENA modules also reveal functional consequence in other functional processes, for example, ‘Morphogenesis’, ‘Signaling Cascade’, ‘Differentiation & Development’ in c1_90 (Supplementary Fig. 7C) and ‘Metabolism and Biosynthesis’, ‘Lipid & Membrane Metabolism’ and ‘Transcription & RNA Regulation’ in c1_247, being their top 3 dominant functions (Supplementary Fig. 7E).

### MEGENA revealed several overshadowed neuronal functional consequences from ADHD-associated co-expression modules in the Putamen

Similar to caudate, putamen’s MEGENA analysis in addition to identifying functional themes consistent with DEGs enrichment analyses, also several neuronal functional themes revealed 16 modules functionally associated with ‘Synaptic Function & Neurotransmission’ as well as with ‘Cellular Adhesion, ECM and BBB integrity’ (Supplementary Fig. 5B), with 3 modules of those modules having these as amongst the top 3 functions (Fig. 2F). Functionally, 8 modules are associated with ‘Neuronal & glial Differentiation & Development’, 11 with ‘Ion Homeostasis and Membrane Excitability’, 12 with ‘Vasculature and Angiogenesis’, 7 with ‘Neuronal connectivity and Circuit Formation’, 6 with ‘Brain Morphogenesis’ and 7 with ‘Other Glial Processes and Functions’(Supplementary Fig. 5B), where several modules have these in their top3 functions (Fig. 2F). However, considering top 3 ranked functional themes as representative/ dominant functions of the modules, ‘Synaptic Function & Neurotransmission’ was a dominant function of 3 modules, ‘Cellular Adhesion, ECM and BBB integrity’ and ‘Ion Homeostasis & Membrane Excitability’ were dominant functions of 3 modules each i.e., 9 modules in total (Fig. 2F). In addition, ‘Neuronal & Glial Differentiation & Development’ which annotates for BPs related to growth of neurons and glial cells and their differentiation through processes such as neurogenesis, gliogenesis, cell migration and maturation etc., and ‘Other Glial Processes and Functions’ which annotates for BPs related to functions of all glial cells such as myelination, neurotransmitter reuptake, etc. (Supplementary Table 18, 16) and ‘Neuronal Connectivity & Circuit Formation’ were also identified to be dominant functions of 1 module each i.e., 3 modules in total (Fig. 2F).

Similar to the caudate, functional specification from parent leaf modules was also observed in the putamen. In the family of c1_110 (parent) and c1_307 (leaf), from the parent’s 24 functional themes, the leaf module shares 14 functional themes (Supplementary Fig. 8), becoming more neurologically related. It also governs several neuronally centered functions, including ‘Neuronal & Glial Differentiation & Development’, ‘Brain Morphogenesis’, ‘Other Glial Processes & Functions’, ‘Cell Adhesion, ECM & Barrier Integrity’ and ‘Intracellular Transport & Trafficking’, with the first one being the module’s one of the dominant functions (Supplementary Fig. 8C). The parent module c1_110 too, was associate to ‘Cell Adhesion, ECM & Barrier Integrity’ as its dominant function as well as with ‘Neuronal & Glial Differentiation & Development’ and ‘Intracellular Transport & Trafficking’ as neuronal non-dominant functions (Supplementary Fig. 8A). The gene specification from parent to child module in this family stands out as it goes from a gene set of 50 genes from parent c1_110 module to a gene set of 14 genes in lead c1_307 module (Supplementary Fig. 8B, D: network plots, Supplementary Table 12: module size; Supplementary Table 15: gene list).

Unlike caudate, the putamen had modules with positive as well as negative correlations to ADHD. Interestingly, modules associated with ADHD in opposite directions were found to share the same dominant functional theme. Modules c1_199 and c1_88 both had ‘Ion Homeostasis & Membrane Excitability’ as their top 1^st^ ranking functional themes (Supplementary Table 18, Fig. 2D, Supplementary Fig. 9A); however, the first has a negative correlation to ADHD, and the latter has a positive correlation to ADHD (Supplementary Table 12). Owing to the same, some parent-leaf module pairs too shared opposite correlation with ADHD, unlike the earlier mentioned c1_110 (parent) and c1_307 (leaf) both are negatively correlated to ADHD, for instance, c1_19 and c1_113 where the earlier is negatively correlated and the later is positively correlated to ADHD phenotype.

Several putamen’s modules showed association with more than 1 neuronal functional theme, such as c1_110, c1_307, c1_186, and c1_99, to name a few, but none had more than 1 neuronal functional theme in its top 3 dominant functional themes, unlike caudate’s module c1_256. However, module c1_199 was particularly interesting as, along with being neuronally centered, its top dominant functional theme, ‘Ion Homeostasis & Membrane Excitability’, has a rank of >50 (Supplementary Table 18, Fig. 2D). The module comprised of 10 genes with KCNE1 as the regulatory hub gene of the network, which encodes a subunit for assembling potassium voltage-gated channel. In all, 12 modules were found that had neuronal functional theme as one of its top dominant functions, such as positively correlated module c1_390 has ‘Synaptic Function & Neurotransmission’ as its top dominant function and negatively correlated module c1_12 has ‘Neuronal Connectivity & Circuit Formation’ as its top dominant function (Supplementary Table 18).

In addition to modules governing several neuronal processes, putamen’s MEGENA modules also revealed functional consequences in other functional processes, for example, ‘Cytoskeleton & Structural Dynamics’ and ‘Signaling Cascade’ in c1_186 (Supplementary Fig. 9C) and ‘Lipid & Membrane Metabolism’ and ‘Cellular Homeostasis’ in c1_99, as being amongst the top 3 dominant functions of these modules along with ‘Immune & Inflammatory Processes’ (Supplementary Fig. 9D).

### Metabolomics Implicate Dopamine as Top Differentially Expressed Metabolite in ADHD

Differentially expressed metabolites (DEMs) within the Putamen and Caudate were identified through metabolomics of ADHD and Control cohorts. Several distinct metabolites were identified in each tissue type at a threshold of p-value <=0.05, including Citrulline (Cit) within the Caudate, as well as Dopamine, Succinic acid (Suc), and Phosphatidylcholine 30:0 (PC 30:0) within the Putamen (Fig. 3A & B, Supplementary Table 19 & 20). A pooled analysis combining ADHD and Control samples across the caudate and putamen cohorts was also performed. Dopamine, succinate (Suc), and citrate (Cit) remained significantly altered in this pooled analysis, along with arginine (Arg) and asparagine (Asn), which also emerged in the pooled analysis (Fig. 3C, Supplementary Table 21). Comparison of all six DEMs identified in the analyses were examined across pooled Putamen and Caudate samples, where bidirectional agreement in fold-change was observed for all DEMs except PC 30:0 (Fig. 3D). Per-metabolite scaling was then applied to determine relative DEM enrichment between ADHD and Controls, where a stark within-group contrast was observed for Dopamine, Suc, Arg, Asn and PC 30:0 between ADHD and Control cohorts of putamen (Fig. 3E), and for Cit, Asn and Dopamine between ADHD and Control cohorts of caudate (Fig. 3E). Throughout the analyses, Cit and PC 30:0 were the only DEMs with higher relative abundance compared to Controls in the ADHD cohort (Fig. 3F). Pooled ADHD and Control DEMs displayed trends similar to those within each individual cohort type (Fig. 3F, G).

**Figure 3:**
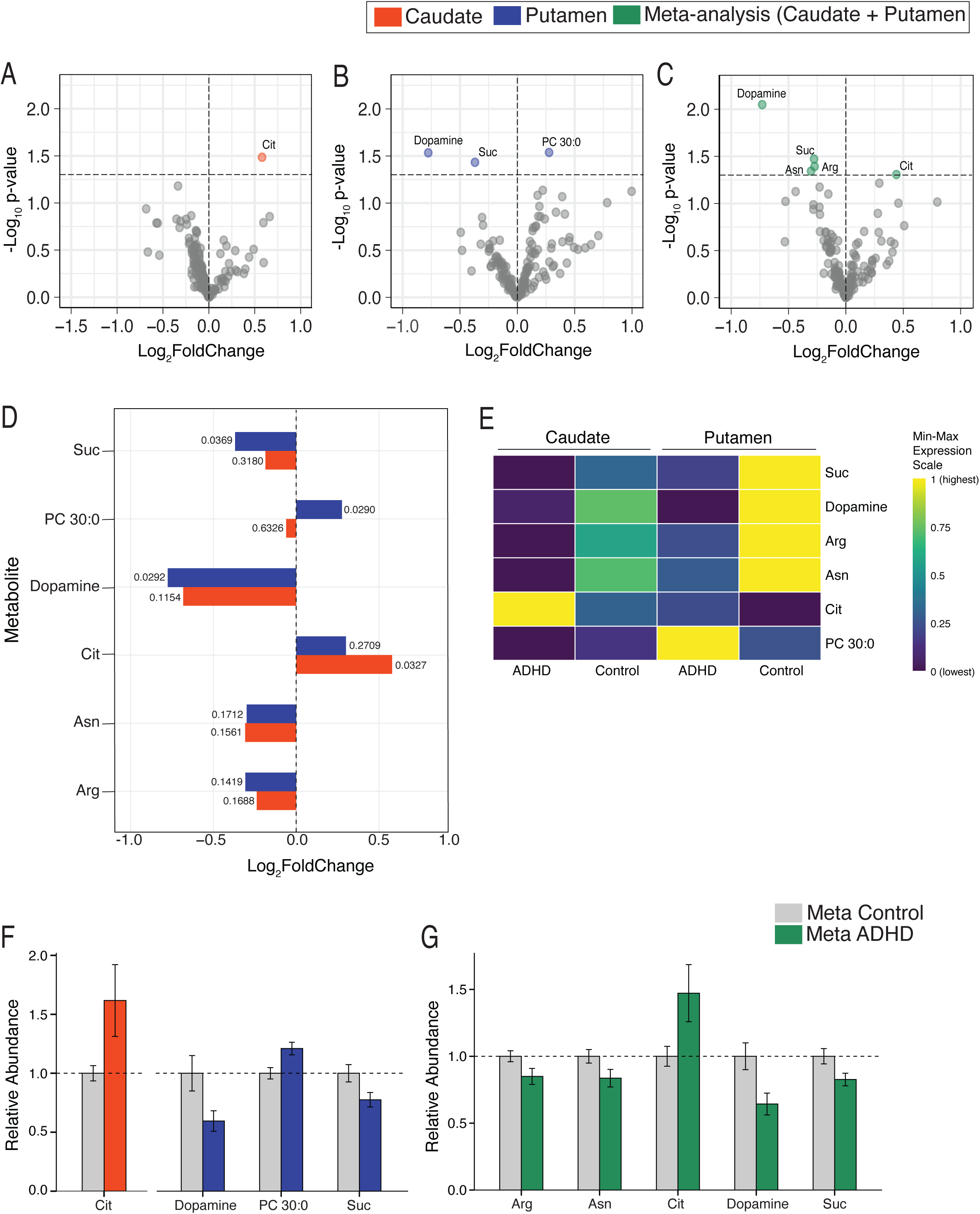
Metabolome of Caudate and Putamen in ADHD: (A-C) Volcano plots identifying the differentially expressed metabolites (DEMs) in the ADHD group as compared to Control group individuals within the Caudate (A), the Putamen (B), and in pooled Caudate and Putamen samples (C). The significant DEMs, i.e., the metabolites with p-value =< 0.05 (marked with a dotted horizontal line), are represented in orange for Caudate, blue for Putamen, and green for the pooled analysis. (D) Bar graph exhibiting all the significant DEMs across Caudate, Putamen, and Meta-analyzed datasets and their expression (log2 fold change on x-axis) in Caudate and Putamen samples individually, with the corresponding region-specific p-values mentioned beside each bar. (E) Heatmap of DEMs showing expression scaled by metabolite in Control versus ADHD groups in Caudate and Putamen tissues. Expression is scaled independently for each metabolite, such that for a given metabolite the lowest group mean amongst the 4 groups is at 0 and the highest is at 1. (F, G) Relative abundance of each significant DEM in the ADHD group as compared to the control group. Relative abundance was calculated by setting the Control group to 1 (marked by horizontal dotted lines); bars represent the group means and error bars represent the standard error of the mean. The Panel F bar graph highlights metabolites significantly differentially expressed in the Caudate and Putamen individually, and panel G highlights those identified in the pooled analysis for ADHD and Controls. In total, 162 metabolites were analyzed using Liquid chromatography–mass spectrometry (LC-MS). Caudate and Putamen tissue was obtained from the same 14 individuals (Control = 7, ADHD = 7), giving 14 samples per brain region and 28 samples in total.

### Cross-study comparison analysis identifies genes emerging across studies in caudate as well as putamen brain regions

Lastly, we did a comparative analysis our results with the 2 existing studies, Sudre et al.^43^ study the only bulk RNA-sequencing study in ADHD to be conducted on post-mortem brain tissue till date and Demontis et al.^12^, one of the biggest ADHD GWAS study till date, which identified 27 genome-wide-significant ADHD risk loci mapped to 76 plausible risk genes.

Upon comparing significantly dysregulated genes in caudate brain region between our study and Sudre et al. study, 35 genes were found to be dysregulated in both studies, and 29 had a fold change of >= |2| (Fig. 4A, Supplementary Table 22,) i.e, were HES-DEGs in both. Surprisingly, 32 of these had dysregulation in opposite directions in both studies, where most, i.e., 29 genes, demonstrated upregulation in our data and downregulation in Sudre et al.’s (Fig. 4A, Supplementary Table 22). In addition to the dysregulation in opposite directions, the magnitude of fold-change observed in our study, for most genes, was much larger than reported in Sudre et al.’s study in the opposite direction (Fig. 4A).

**Figure 4:**
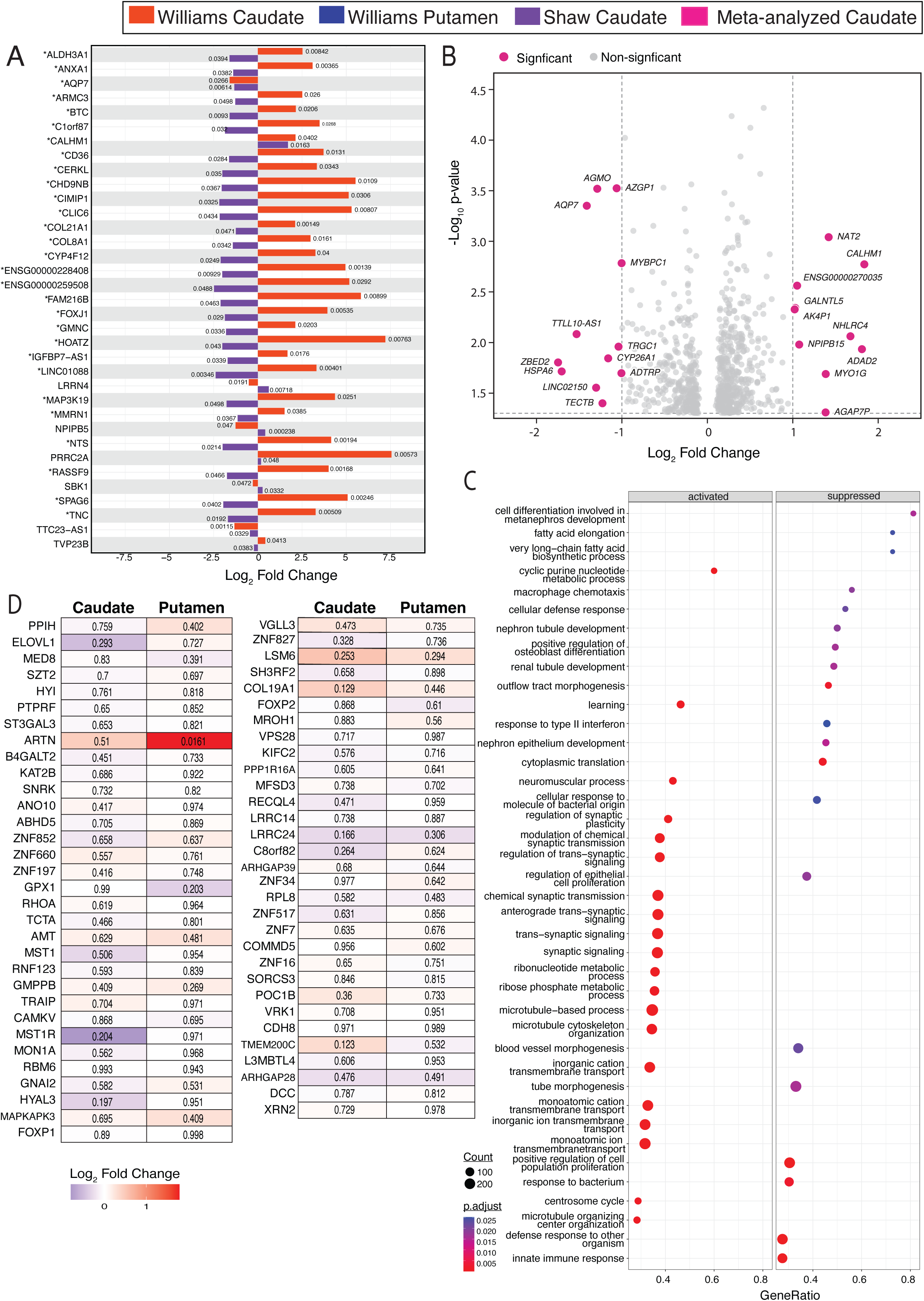
Comparative analysis of our Bulk RNA-sequencing data with exisitng large-scale studies: (A) Bar graph of significant DEGs overlapping between our and Shaw groups’ (i.e., Sudre et al.) Caudate Bulk RNA-sequencing datatsets and representing their expression (Log2Fold Change on x-axis) in both datastes individually with the p-value mentioned besides the correpsonding bars. The DEGs marked with * are significant and have fold change >= 2 (i.e., are Highly Expressed Significant DEGs (HES-DEGs)). (B) Volcano plot representing the meta-analysis of Caudate Bulk RNA-sequencing from our and Shaw group’s datasets. The colored dots represent HES-DEGs with the corresponding gene names labelled besides each dot, and the thresholds (p-value =< 0.05 and fold-chnage >=2) are marked by dotted lines. (C) Biological pathways associated with the cumulative effect of significnat DEGs identified in Meta-analyzed dataset using GSEA analysis, along with the direction of their effect i.e. in up-(activated) or down-(suppressed) regulated manners. (E) Gene expression of 76 potential genes mapped from 27 genome wide significant ADHD risk loci as identified by Demontis et. al., in our Caudate and Putamen tissues’ DEGs. 63 out of 76 genes were present in our identified Caudate and Putamen’s DEGs, p-value of these DEGs is mentioned inside each gene’s boxes in the heatmap and their Log2FoldChange is represented by the color of the box on the scale mentioned in the figure, where red indicates a postive fold change and blue indicates a negative fold chnage. (Significant DEGs are the ones with p-value =< 0.05).

To further evaluate these two studies methodically to integrate evidence across both studies and assess between-study heterogeneity, meta-analysis of these studies was performed. Most of the genes with meta-analysis estimates had I^2^ between 0-3% (Supplementary Fig. 12A, Supplementary Table 23). The meta-analysis identified approximately 809 significant DEGs with 23 HES-DEGs (Fig. 4B, Supplementary Table 23). 491 of the 809 genes exhibited I^2^ between 0% to 11% (Supplementary Fig. 12C, Supplementary Table 23). But to characterize cross-study consistency, these 809 DEGs were evaluated based on not only I² (of these in the meta-analysis), but also on the effect direction (i.e. direction of fold-change in their respective bulk RNA-sequencing analyses). 470 genes showed concordant directions of effect across both studies with I² ≤ 50%, indicating low-to-moderate estimated heterogeneity (Supplementary Fig. 12D, Supplementary Table 23). Notably, 409 of these 470 genes had I² between 0% to 5%, indicating very low estimated between-study heterogeneity (Supplementary Fig. 12E, Supplementary Table 23). On the other hand, 24 of the 35 overlapping DEGs between these two studies, were found to have I^2^ ranging from 69% to 92% and the 9 had I^2^ between 92% to 93.8%, in the meta-analysis (Supplementary Fig. 12B, Supplementary Table 23). This high heterogeneity aligned with the striking difference in the fold-change direction when comparing their values from individual studies.

The GSEA-BP analysis of the 809 significant DEGs from meta-analysis exhibited activation of biological processes related to synaptic signaling, synaptic plasticity, microtubule organization, transmembrane transport and a few metabolic processes (Fig. 4D, Supplementary Table 24). In contrast, suppressed biological processes were related to immune response, cell differentiation and proliferation, morphogenesis and development of tissues and organ systems, and cytoplasmic translation (Fig. 4D, Supplementary Table 24). The top 5 BPs, were all related to synaptic signaling, including modulation of chemical synaptic transmission and regulation of trans-synaptic signaling (Supplementary Fig. 11).

Upon conducting a stringent three-way overlap analysis of significant DEGs both studies independently and from the meta-analysis, only 5 DEGs result, namely AQP7, CALHM1, PRRC2A, NPIPB5 and TTC23-AS, with the first two being the HES-DEGs across all 3 analyses (Supplementary Fig. 10A & B). Whereas 42 genes overlapped between our study and meta-analysis’ DEGs (Supplementary Table 25, Fig. 10C) and 8 of those genes (MYO1G, ADAD2, AQP7, CALHM1, TTLL10-AS1, NHLRC4, AK4P1 and AGAP7P) were HES-DEGs (Supplementary Fig. 10D).

Lastly, the expression levels of the 76 potential risk genes mapped by Demontis et al., from 27 genome wide significant loci were analyzed. Out of 76, 63 genes were found amongst our caudate’s and putamen’s DEGs dataset However, none of those genes in caudate brain region had significantly dysregulated gene expression. In contrast, ARTN in the putamen was identified to have significantly dysregulated expression, with a fold change of 3.46 (Fig. 4E, Supplementary Table 26).

## Materials and methods

### Cohort design and sample acquisition

Post-mortem tissue from 2 brain regions, namely caudate and putamen, were acquired from the National Institutes of Health biobank and Human Brain Collection core. Several criteria were laid out for sample selection, before sample acquisition. The maximum age limit was set to 18 years. The study also focused on one particular ADHD subtype: the combined type, to reduce variability. All case (i.e., ADHD) samples were also required to have a clinical diagnosis of the phenotype, with no cases having schizophrenia as comorbidity. This study was kept inclusive of both sexes and so included samples from both male and female individuals. However, owing to low sample availability at the biobanks that would match our stringent criteria, the age limit was extended to 19 years, and 6 samples above 18 years were included to increase sample size, without substantially over-ruling our criteria. The samples thus acquired were between the ages of 12 and 19 years. 28 samples matched these criteria and the final cohorts consisted of 14 samples per brain region, each comprising of 7 ADHD and 7 control samples. All individuals/ donors comprosomg ADHD cohort had a prior clinical diagnosis of ADHD from a clinician, five of which included a comorbid diagnosis that included mood disorders such as bipolar disorder, major depressive disorder, or post-traumatic stress disorder (Supplementary Table 1). No comorbidities were reported in controls (Supplementary Table 1). All 14 caudate and putamen tissue samples were obtained from the same donors across both brain regions to minimize variability. Detailed information on each sample can be found in Supplementary Table 1.

### RNA extraction and sequencing

The process from RNA extraction to sequencing was conducted by Genomics and Bioinformatics Core of University at Buffalo. RNA was extracted from <=50mg of tissue using the TruSeq Stranded RNA kit. RNA concentration was quantified, and RNA quality was assessed using the Agilent Bioanalyzer. Based on the core facility’s quality-control (QC) assessment, 18 samples passed QC and were selected for sequencing, comprising 9 samples per brain region, with 6 ADHD and 3 control samples in each region. Among these nine samples per brain region, eight caudate–putamen pairs were acquired from the same individuals/ donors, while one sample in each region came from a different individual/ donor (93052 in caudate’s and 93027 in putamen’s control cohort).

Paired-end RNA-sequencing was conducted for all 18 samples on NovaSeq 6000 using an SP flow cell with a read length of 2 × 100 bp. Following sequencing, reads were processed with fastp for quality filtering. Post-processing quality was assessed with FastQC and summarized across samples using MultiQC; all samples passed QC and were retained for downstream analysis. The reads were aligned using STAR^52^ and indexed against GRCh38 reference genome as made available by GENCODE release 21^53^. Gene-level read counts were generated using featureCounts from the Subread package, and genes with fewer than 10 counts in at least three samples were excluded prior to differential expression analysis. Differential expression analysis was then performed using DESeq2 package^54^ form Bioconductor to identify genes that are differentially expressed in ADHD cohort as compared to control group. This identified 20,009 DEGs in caudate and 20,260 in putamen, of which 432 in caudate and 347 in putamen passed the statistical significance determined using the Wald test with *p*-value ≤ 0.05 were considered significantly differentially expressed genes. Ensembl gene IDs were mapped to HGNC gene symbols using biomaRt^55, 56^ R package with the Ensembl hsapiens_gene_ensembl dataset, to convert them to gene symbols. In instances where an Ensembl ID did not have a corresponding gene name, the gene’s Ensembl ID was retained. The volcano plot was plotted using ggplot R package and significantly differentially expressed genes were marked with thresholds of p-value <= 0.05 and a fold change of >= |2| i.e., log2FoldChange >= |1|.

### Pathway and function enrichment analysis of DEGs

Gene set enrichment analysis^44^ (GSEA) was performed independently for the caudate and putamen using the gseGO() function in the clusterProfiler R package^45^. Genes were ranked according to the DESeq2’s Wald test statistic (stat) and analyzed against Gene Ontology (GO) Biological Process (BP) and Molecular Function (MF) independently. Human gene annotations were obtained from Bioconductor org.Hs.eg.db, with Ensembl gene identifiers used as input. Gene setsl containing 10–800 genes were included in the analysis, and enriched terms were retained using a p-value cutoff of 0.05 with 100,000 simple permutations.

Enrichment results were visualized using dot plots and gene–concept network plots. Dot plots displayed up to the top 20 positively enriched (activated) and top 20 negatively enriched (suppressed) BP or MF terms, with dot size representing the number of terms present for that theme, dot color representing the adjusted p-value and the x-axis representing the Gene Ratio. Gene Ratio is defined as the proportion of genes contributing to the enrichment of a given BP or MF term relative to the total number of genes associated with that GO term. Gene–concept network plots were generated for the top five enriched GO terms, irrespective of enrichment direction, to visualize the input genes contributing to each enriched GO term.

### Multiscale Embedded Gene Co-expression Network Analysis (MEGENA)

#### Co-expressed gene networks (modules) identification

Raw gene count data generated using featureCounts was filtered to retain genes with >=10 counts in at least three samples. From the filtered count matrix, the top 2500 DEGs (ranked by (-log10p-value) × absolute(log2fold-change)) as identified from the earlier DESeq2 analysis identifying DEGs were taken further for gene co-expression analysis to identify modules governed by the most differentially expressed genes. Gene expression counts were variance-stabilized using DESeq2 package, and the transformed expression matrix was used for MEGENA network construction. It was implemented using the MEGENA R package^50, 57, 58^ and according to the framework described by Song and Zhang^50^, except, we used top ranked 2500 DEGs. Gene–gene Pearson correlations were calculated using a top-K approach (K = 300), retaining strong correlations irrespective of direction (use_abs = TRUE), followed by planar filtered network (PFN) construction. Co-expression modules with a minimum module size of 10 genes and maximum size of 2500 genes were set to be identified using the MEGENA R package^50^. This was performed for caudate and putamen independently, and 177 and 187 modules were identified for caudate and putamen, respectively. Following module identification, for each MEGENA module, the module eigengene (ME) and eigengene-based connectivity (kME) were calculated using WGCNA functions^57, 58^. ME is the first principal component of the expression matrix of genes within that module and representing the summary expression pattern of the modules, and kME indicates which genes are most representative within the module. Lastly, the correlation of each module with ADHD (i.e., r) was also assessed by computing the correlation coefficient (r) between each module’s ME and the trait. The modules significantly correlated with ADHD i.e., with p-value <= 0.05, were taken further for module pathway enrichment analysis.

### Pathway enrichment analysis and functional annotation of significantly correlated modules

The functional enrichment analysis was performed programmatically by accessing the EnrichR^46, 47^ application programming interface (API) using enrichR() R package^51^, with Gene Ontology Biological Process^48^ (GO-BP) library (version 2025). This enabled automated submission of gene lists from caudate and putamen’s significantly correlated modules and retrieval of their enrichment results. However, as ontology analysis usually gives a long list of all associated GO terms associated with the input gene list, which can be both redundant and difficult to interpret, we curated a list of 30 broad functional themes. We defined each theme, with functions and biological processes that would fall under them along with few examples of GO terms that would be classified under them. The biological processes that usually function together or have overlapping pathways were combined, such as neurotransmitter availability at the synapse, which is closely related to synaptic function, were combined as Synaptic Function and Neurotransmission, including calcium ion (Ca^++^) movement, as that is closely related to neurotransmitter release at so ita availability at the synaptic cleft. However, keeping these themes distinct enough that membrane excitability was defined as a separate theme, with Na^+^, K^+^, and Cl^-^ movements categorized under Ion Homeostasis and Membrane Excitability, as their functions are closely related to the action potentials (membrane excitability). So themes were curated to keep biological functions that share biological processes under the same theme but distinct enough to identify specific functions distinctly e.g., proteostasis can be analyzed via 2 distinct themes: one related to its synthesis and processing i.e., “Protein Synthesis & Processing” and the other related to its quality detection, response to its dysregulation, and its degradation, i.e., “Protein QC, Stress & Degradation”. Thus, the resulting final list of 29 functional themes (Supplementary Table 16) was curated to be non-overlapping, specific, and interpretable.

Once the functional themes were curated and defined, GO-BP enrichment results for each module were classified, with each GO term in GO-BP results being classified under particular functional themes. Next, for each module, we ranked all its functional themes by a ranking metric that we created, incorporating the number of GO terms in the theme (n), the top GO term in the theme’s (-log_10_p-value), and its combined score as obtained from the enrichment analysis normalized on the scale of 0 to 1, culminating in:

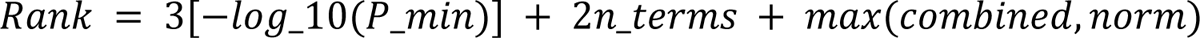

As the top 3 functional themes captured the largest proportion of GO terms associated with the module, for most modules, the top 3 functional themes were assigned as the module’s (dominant) functions, and hence the module’s annotated functions. We also consolidated a dictionary of 1,909 GO terms (identified across all modules’ enrichment results) and their classification under their functional theme, for future use and reproducibility purposes.

### Data Visualization of MEGENA results

For data visualization, interpretation, and analysis purposes, several graphs were generated. A heatmap was plotted with the MEs of every sample (ADHD and control) for all significantly correlated modules. Hierarchal plots were generated to visualize the relation between all modules starting from root module with branchings into smaller parent modules to their end leaf modules. Network plots were generated for every module, demonstrating the connectivity between all genes in the module and highlighting the hub gene of the module. Excel sheets with every module’s gene list (compiled in Supplementary Table 14 and 15) and Excel sheets with a summary file compiling all modules with their r, p-value, module type, size, hub genes, significance, and more (Supplementary Table 11 and 12).

Next, from the enrichment and functional annotation results, we generated dot plots for each module exhibiting its top 10 functions, with dot size corresponding to the number of GO terms classified for the theme (n). Two heatmaps were generated, one exhibiting every modules’ top 3 functional themes along with a count of total number of modules that have the functional theme as its (a) top 3 function i.e. dominant function and (b) as its function irrespective of rank. The other heatmap exhibited all modules with all its functions and number of GO terms classified in every functional theme for every module represented in the tiles of the heatmap.

### Cross-study comparative/ meta-analysis with studies in existing literature and within our study between the two brain regions

#### Shared DEGs between 2 studies in caudate

RNA-Seq data generated from our caudate brain tissue samples was meta-analyzed with bulk RNA-seq data from the previously published Sudre et al.’s^43^ “Mapping the cortico-striatal transcriptome in attention deficit hyperactivity disorder” study. Data from this study was generated from anterior cingulate cortex (ACC) and caudate nucleus brain regions in individuals diagnosed with ADHD through clinician interviews with next of kin. Bulk RNA-seq data were downloaded from their supplementary file 2, which uses the largest subpopulation (white, non-Hispanic) in their study, as it matches our study’s largest subpopulation (Supplementary Table 1).

To match the format of our data, ensembl IDs in Sudre et al.’s dataset were converted to HGNC gene symbols using biomaRt R package^55, 56^ and Ensembl hsapiens_gene_ensembl dataset, as described earlier. For instances where an ensembl ID did not have a corresponding gene name, the gene’s ensembl ID was retained. To ensure compatibility, data from our caudate samples, our putamen samples, and Shaw’s caudate samples were combined into a single master table file, containing Ensembl IDs, gene symbols, log_2_Fold Change, and p-value from all 3 datasets. DEGs that passed the threshold of p-value < 0.05 in both, our caudate samples and in Sudre et al.’s caudate samples i.e., the significant DEGs across both studies, were plotted using ggplot package in R for data visualization and analysis. The significant DEGs, that passed the threshold of log_2_Fold Change >= |2|, were deemed highly-expressed significant DEGs (HES-DEGs) and were marked with * in the graph.

#### Shared DEGs between caudate and putamen

The same workflow as above was used for identification of shared DEGs between our caudate and putamen datasets, i.e., by using the master table file, identifying the shared significant and HES-DEGs using the above-mentioned thresholds, and plotting the results using the ggplot R package with HES-DEGs marked with *. Subsequently, Gene Ontology analyses^48^ for Biological Process, Molecular Function and Cellular Component as well as Synaptic Gene Ontology^49^ enrichment were conducted on the overlapping up- and down-regulated HES-DEGs independently, using EnrichR’s web-based platform^46, 47^.

#### Meta-analysis of two caudate bulk RNA-sequencing analyses

The DEGs file from Sudre et al.’s study’s caudate was acquired from their Supplementary File 2 to meta-analyze with our study’s caudate’s DEGs. For this, firstly, weights were defined for both studies based on their standard error (SE), to take into account the variability between studies and to consider in all subsequent calculations. Following this, all statistics were calculated from both studies for the meta-analysis including (a) effect size of all genes from both studies combined i.e., the weighted average of log_2_FoldChange, (b) variance of the meta-analysis’ estimates i.e. SE, (c) Z, (d) p-value for each gene and lastly running a heterogeneity diagnostics for each gene by (e) Cochran’s Q and (f) I^2^, reflecting the heterogeneity between the studies. A volcano plot was also plotted for visualization of significant DEGs with fold change >= |2|. To test heterogeneity between these two bulk RNA-sequencing studies, several histograms of I^2^ were plotted for different gene sets namely (i) DEGs overlapping between the 2 across bulk RNA-seq analyses studies, (ii) all genes with meta-analysis estimates, (iii) significant DEGs identified by metanalysis, (iv & v) from these DEGs and HES-DEGs the one with I2 =<50% and have same direction of effect (i.e. fold-change) the 2 studies; by filtering the rows from the meta-analysis file and plotting using Excel functions.

GSEA^44, 45^ pathway enrichment analysis was conducted against GO-BP library and visualization was done through dot plot and network plot as mentioned earlier under ‘Pathway and functional analysis of DEGs’.

#### Shared DEGs across 3 analyses from 2 studies in caudate

Gene-set overlap analyses were conducted. Pairwise intersections were first assessed between our caudate dataset and the meta-analyzed caudate (MC) dataset, followed by a three-way intersection including the Sudre et al. caudate dataset. The analysis was performed using the online Interactive Venn tool^59^, which helped identify the overlapping genes from the input of 3 gene lists. This was conducted first for significant DEGs and then for HES-DEGs.

#### Shared genes between our bulk RNA-sequencing-identified DEGs and previously GWAS-identified potential risk genes

Bulk RNA-seq data from our human adolescent caudate and putamen tissue samples were analyzed against genes surrounding potential ADHD risk loci identified by Demontis et al. in “Genome-wide analyses of ADHD identify 27 risk loci, refine the genetic architecture and implicate several cognitive domains”^12^. This study conducted a large genome-wide association study (GWAS) meta-analysis involving 38,691 individuals with ADHD and 186,843 controls. The study identified 27 genome-wide significant ADHD risk loci that mapped to 76 plausible potential risk genes using positional and functional genomic evidence^12^. We filtered these 76 genes in our caudate and putamen DEGs datasets using the master table file created and plotted a heatmap of their expression using the ggplot2 R package.

### Metabolomic Profiling

All 28 post-mortem tissue samples (14 from caudate and 14 from putamen) were processed were processed through the Biocrates MxP Quant 500 kit (Biocrates Life Sciences AG, Innsbruck, Austria) offered through the Bioanalytics, Metabolomics, and Pharmacokinetics Shared Resource (BMPK) at Roswell Park Comprehensive Cancer Center. The Biocrates MxP Quant 500 panel quantifies 630 metabolites across 26 biochemical classes using liquid chromatography–tandem mass spectrometry (LC-MS/MS). Raw readouts from the panel were imported into R for processing. From the full panel, only those with detectable values in at least 50% of samples were retained. This resulted in a total of 162 metabolites used for downstream analysis. Within the 4,536 remaining measurements (28 samples × 162 metabolites), 153 values were missing; these were imputed as one-half of the lowest quantified concentration observed for that metabolite. Concentrations were next log_2_-transformed to normalize the dataset. Effect sizes between cohorts and tissue types were computed through an Empirical Bayes procedure performed by the limma package^60^ in R. Differential abundance was assessed separately within each brain region (n = 14 per region; 7 ADHD and 7 control) and, in a pooled analysis across all 28 samples together. Metabolites with a nominal *p*-value below 0.05 were considered significant DEMs and were selected for visualization through various cross-cohort analyses.

Volcano plots for DEMs highlighted significant DEMs in colors specific by tissue region. A bar graph of all significant DEMs was plotted illustrating their Log_2_FoldChange observed in caudate and putamen individually for comparing their fold-change between the 2 brain regions. A heatmap was used to scale expression of DEMs in Control and ADHD cohorts for Caudate versus Putamen tissues. Expression was scaled independently for each metabolite, such that for a given metabolite the lowest group mean amongst the 4 groups (2 cohorts per brain region) is at 0 and the highest is at 1. Bar graphs were used to visualize the relative abundance of each significant DEM in the Control versus ADHD cohorts, for Caudate, Putamen, and pooled analyses individually. Relative abundance in the ADHD group was calculated relative to the Control group by normalizing control group metabolite values to 1, with bars representing the cohort means and error bars representing the standard error of the mean.

## Discussion

Along with identifying several overlapping mechanisms between the interconnected brain regions, caudate and putamen, at the transcriptomic level 20 shared DEGs were found, which account for only 2.64% mRNA-level commonality between both regions, indicating that at the transcription level both regions are affected in a considerably different manner. Both regions reported reduction in dopamine levels from pooled/ combined analysis, with considerable reduction in putamen in region-specific metabolomics analysis, along with an increase in calcium ion transport and synaptic organization-related pathways, which can be compensatory mechanisms in response to the reduced dopamine levels. However, several shared biological processes such as ATP synthesis and immune response were dysregulated in opposite direction. Region-specific alterations included developmental delay and increased myelination in the caudate, and increased neurotransmitter reuptake in the putamen. Our overlapping and contrasting findings between caudate and putamen support the existing view that altered brain functions are based on disrupted networks across brain regions rather than an individual brain region^7, 61^, which is reflected from the overlapping pathways between caudate and putamen. But at the same time, the contrasting pathways demonstrate that region-specific alterations in ADHD do occur, and therefore, investigating individual brain regions at genetic, transcriptomic, and metabolomic levels is necessary to obtain a comprehensive understanding of the disorder. In this study, we also highlight multiple dysregulated genes and pathways from several analyses, including transcriptomic, gene co-expression, and cross-study validation analyses. With these findings, our study is the first to report transcriptomic dysregulation in the putamen, metabolomic dysregulation in putamen and caudate, and to report the shared and contrasting pathways affected between the regions that together form the dorsal striatum. It is also the first study reporting age-specific alterations in the dorsal striatum in ADHD.

Dysregulation in dopamine signaling, its related circuitries, and its association with ADHD symptoms has been identified and established from several studies^62, 63, 64, 65^. Our study provides direct evidence and quantification from postmortem tissue, exhibiting a 1.7-fold reduction in dopamine as a metabolite itself in putamen and a reduction of 1.66 times in pooled analysis of dorsal striatum with even greater significance (p-value = 0.00893). In addition, this study also revealed that the reduction in dopamine is more evident in putamen than in caudate. Citrate (Cit), an important intermediate in the tricarboxylic acid (TCA) cycle of cellular respiration^66^, participates in the regulation and metabolism of glutamate and GABA^67, 68, 69^ and lipid synthesis^70^, was the only metabolite with increased levels, particularly in caudate and pooled dorsal striatum, whereas PC 30:0, a phosphatidylcholine and a component of cellular membranes, was increased only in putamen. On the other hand, succinate (Suc), which is also an intermediate of TCA cycle^71^ and plays a role in metabolic and immune signaling^72^; asparagine (Asn), which participates in biosynthesis and brain developmen^73, 74^ and arginine (Arg), which also participates in cellular metabolism processes in the nervous system^75, 76, 77^ were all reduced in pooled dorsal striatum analysis with succinate also showing reduction in putamen. Notably, of all the identified metabolites (DEMs), dopamine exhibited the highest altered expression levels with fold change of 1.7 and 1.66 times.

The levels of all metabolites, given their biological functions, substantiate the dysregulated biological processes and mechanisms revealed from the significant DEGs, gene set enrichment, and co-expressed gene networks (i.e., MEGENA) analyses. For instance, in line with the roles of the above metabolites, MEGENA identified several caudate and putamen modules reported signaling pathways, immune and inflammatory processes, lipid-, membrane- and energy-metabolism, biosynthesis, and proteostasis as their dominant functions; and DEGs enrichment analysis reported strong perturbation in mitochondrial ATP synthesis biological processes, concordant with altered levels of Cit and Suc and their role in TCA cycle i.e., cellular respiration.

Although only 13 HES-DEGs were shared (out of 517 across both brain regions), the up-regulated HES-DEGs governed processes and functions that influence neurotransmitter signaling/availability. GO and SynGO enrichment analyses via GSEA revealed that the EFHB gene, governed processes related to calcium ion movement via store-operated calcium entry and calcium-dependent calcineurin-NFAT signaling cascade, and molecular functions related to calcium-ion sensing. EFHB is a cytosolic calcium sensor; when it detects endoplasmic reticulum calcium store depletion, it activates downstream components promoting STIM1-Orai channel coupling, allowing calcium to enter the cell^78, 79^. Sustained calcium influx in turn causes activation of calcineurin-NFAT signaling, which supports axon growth, projection, synaptic remodeling as well as neural development^80, 81^. These findings, were complementary to SynGO’s identification of increased postsynaptic organization and modulation of synaptic transmission by up-regulation of ROR2 gene. Together, these suggest that shared upregulated genes influence processes that may in turn be the compensatory responses of the ADHD brain, to reduced dopamine levels in both these regions. Additionally, increased levels of Cit in caudate and its potential relevance to glutamatergic and GABAergic neurotransmission^66, 68, 69^; along with the MEGENA revealing modules with dominant influence on synaptic function & neurotransmission (which covers biological processes from neurotransmitter production, transport, release to receptor activity), ion homeostasis and membrane excitability and neuronal connectivity and circuit formation, all implicate dysregulation in neurotransmitter availability at the synapses.

On the other hand, enrichment analysis of the shared down-regulated HES-DEGs was dominated by 3 genes. HSPA1B governing protein stress and QC pathways, metabolic processes of mRNA, ATP and ubiquitin-dependent protein, and regulation of cell growth and survival. HLA-A gene regulating several biological processes involve in immune response and C2CD48 promoting inflammatory response. The reduced expression of these genes indicates down-regulation of these processes. Their cellular component enrichment analysis identified down-regulation of intracellular membrane trafficking via ER–Golgi transport vesicles, endocytic vesicles, recycling endosomes; immune-related cellular machinery via phagocytic vesicles and MHC protein complexes and protein quality-control-structure such as aggresomes.

Immune and inflammatory response dysregulation was also observed from gene-set enrichment analysis of both caudate and putamen DEG gene sets, influenced by multiple DEGs. This dysregulation was particularly stark in the putamen. Interestingly, these pathways were observed to be suppressed in putamen but activated in caudate. Similarly, converse alterations in mitochondrial ATP synthesis processes were also observed, with their activation in putamen, while suppression in caudate. These contrasting alterations in the same biological processes, highlight differential affectedness of caudate and putamen, while also implicating plausible compensatory accommodation in one brain region in response to alterations in the other.

Striatum along with PFC, VTA and NAc is amongst dopaminergic regions in the CNS that have highest levels of dopamine^82^. Dopamine, being an immune modulator, is tightly linked to immune and inflammatory processes^83^. Along with its canonical functions as neurotransmitter involved in cognition, reward, attention and movement, emerging evidence highlights dopaminergic immunomodulation revealing role of dopamine in modulating several immune functions including cytokine secretion, immune cell proliferation, cell adhesion, chemotaxis, antigen presentation and phagocytosis^82, 84, 85, 86^. Most immune cell types express dopamine receptors and dopaminergic proteins and can take up, produce, store and/ or release dopamine^82, 87, 88, 89^. Microglia in humans have been identified to present all 5 dopamine receptor subtypes^88, 89^, however a comprehensive understanding of the dopaminergic system associated with human microglia, still remains to be explored/ understood^82^. This literature provides strong evidence suggesting that the immune and inflammatory alterations observed in the caudate and, more prominently, the putamen may be linked to changes in dopamine levels through underlying cellular mechanisms. Additionally, dopamine’s effect on immune and inflammatory processes is context-dependent, such that it changes depending on its concentration, receptor subtype, cellular activation state and immune cell type^83^. Notably, metabolomic analysis showed dopamine alterations in the same direction in both regions, whereas immune and inflammatory processes were dysregulated in opposite directions. This divergence may reflect region-specific differences in cellular composition, dopamine receptor expression, and/or cellular activation states, which could produce distinct downstream immune responses despite similar directional changes in dopamine levels. Together, these findings further support the hypothesis that the caudate and putamen are differentially affected in ADHD.

ATP synthesis is important for several crucial neuronal and cellular functions, including maintaining Na^+^ and K^+^ ion gradients, especially supporting sustained or prolonged neuronal firing of action potentials^90^. Brain has exceptional energy requirements, being the highest energy-consuming organ of the human body, and synaptic transmission is one of the major contributing factors.^91^ Major energy consumption occurs in synaptic vesicle re-acidification, which helps in refilling of neurotransmitters into the vesicles^92^. Energy is also utilized for postsynaptic ion currents, neurotransmitter recycling, and presynaptic calcium fluxes. A smaller, but non-negotiable fraction of brain ATP goes toward cellular housekeeping including axonal transport, protein and lipid synthesis, cytoskeletal remodeling and immune surveillance and maintenance^93^. According to Faria-Pereira and Morais (2022) synaptic transmission accounts for 43% of grey matter energy demand (i.e., the area where caudate and putamen are located), while maintenance of both neuronal and glial resting potentials represented 15%, action potential propagation represented 17%, and the remaining 25% was assumed for housekeeping processes (including non-signaling processes, such as organelle transport, macromolecules synthesis and degradation)^91, 92^. In fact, energy at the synapse is so crucial that mitochondria are positioned at dendritic ends near synapses^94^ and postsynaptic mitochondria are critical for the development, plasticity, and maintenance of synaptic inputs^95^. The converse alterations in ATP synthesis pathways in the two brain regions not only highlight the difference in affectedness of them, but also more importantly emphasis the dysregulation in energy production in the most energy consuming organ of the body. This alteration is connected to several other alterations observed, especially alterations in synaptic transmission and neurotransmitter availability.

Some interesting DEGs identified in putamen included GABBR1, which encodes GABA_B_ 1 subunit of GABA_B_ receptor that binds to the main inhibitory neurotransmitter gamma-aminobutyric acid (GABA) and mediates slow, long-lasting inhibitory synaptic signal^96^. It showed a 77-fold increase in gene expression (p-value: 0.0024). MOG gene, which encodes the myelin oligodendrocyte glycoprotein, forms the outermost surface of myelin and helps in completion and maintenance of myelin and mediates cell-cell adhesion^97^, showed 1295-fold increased gene expression (p-value: 1.99 E-06). EFHB, as mentioned earlier, mediates axon growth, synaptic remodeling and neuronal development, was increased by 2.71 folds. Additionally, TREM that encodes a cell-surface immunoreceptor protein, predominantly expressed on microglia in the brain^98, 99^. It drives activation of microglia, phagocytosis of. Apoptotic neurons, myelin debris, regulates synapse elimination^99^ and plays an important role in neurodegeneration and neuroinflammation^98^, was down-regulated with a reduced gene expression by 0.489 times (p-value: 0.01).

DEGs identified from the putamen showed predominant suppression of immune and inflammatory processes, including activation, migration and differentitation of immune cells, including microglia activation, as a cumulative effect of gene sets, rather than singular gene functions. Interestingly, it also showed suppression of biological processes inhibiting neurogenesis and causing negative regulation of nervous system development, along with suppression of gliogenesis, neuron remodeling, negative regulation of nervous system, development, neuronal homeostasis, learning, synaptic pruning, regulation of synaptic vesicle exocytosis, regulation of protein modification, response to unfolded proteins, positive regulation of proteolysis, cell adhesion and cytoplasmic translation. On the other hand, putamen gene sets also revealed activation of neurotransmitter reuptake, morphogenesis, nitric oxide synthase biosynthesis, and microtubule bundle formation, in addition to the dominant activation of ATP synthesis processes. These processes reflect the cumulative effects of DEG genes rather than single DEG functions and therefore suggest a more well-rounded and plausible downstream dysregulation resulting from overall transcriptomic perturbations.

Of these biological processes, suppression in regulation of **synaptic vesicle exocytosis, synaptic pruning,** learning, and increased neurotransmitter reuptake collectively emphasize the perturbations in synaptic transmission and neurotransmitter availability at the synapse.

Reduced inhibitions of neurogenesis and nervous system development and increased morphogenesis processes suggest altered regulation of (neuronal) developmental processes, potentially reflecting the well-established implication of developmental delay associated with neurodevelopmental disorders, including ADHD. The down-regulation of biological processes associated with altered synaptic transmission and neurotransmitter availability is also echoed by the increased levels of the inhibitory neurotransmitter receptor GABA-B1, leading to potentially increased inhibition in the putamen and reduced levels of dopamine, respectively. Additionally, reduced neuronal homeostasis reflects reduced arginine levels, and increased ATP synthesis reflects altered citrate and succinate levels, laying a connection between BPs identified from gene set enrichment analysis and metabolomic alterations.

Some interesting DEGs with significantly altered levels include EFHB, as stated earlier; one of the shared DEGs between both these regions was up-regulated by 4.67-fold (p-value: 0.000414), 2-fold more than in the putamen. WNT9A, belongs to the WNT gene family, activates canonical Wnt/β-catenin pathway which further influences cell fate determination and morphogenesis, directs bone and joint development and craniofacial and palatal morphogenesis^100, 101^ was increased by 10.69-fold (p-value: 0.00278), and CALHM1 encodes for voltage-gated and calcium-regulated nonselective ion channel that forms wide pore permeable to large molecules like ATP as well as calcium ions^102^ with a 4.36-fold increase (p-value: 0.000398). Interestingly, amongst the 432 DEGs identified in the caudate, 346 were upregulated, indicating a predominant increase in overall gene expression in the caudate associated with ADHD. This observation warrants further investigation of potential upstream regulatory mechanisms, including enhancers and activating transcription factors, that may contribute to the coordinated upregulation of multiple genes. Identifying such regulatory mechanisms could provide deeper insights into the underlying causes of the observed transcriptional alterations and reveal additional brain-region-specific differences between the caudate and putamen, as the latter exhibited approximately equal proportions of upregulated and downregulated DEGs.

Concordant with the largest proportion of DEGs being upregulated, the gene set enrichment analysis also revealed that most biological processes were activated. However, amongst the few suppressed processes were the CNS myelination, cytoplasmic translation, ribosomal small subunit assembly and ATP synthesis (as described earlier). Despite caudate predominantly being a gray-matter structure, reduced myelination may still reflect altered oligodendrocyte abundance or function and disruption of oligodendrocyte-mediated myelin-associated processes. Amongst the activated processes, firstly, several morphogenesis- and development-related processes, establishment of planar polarity, and the Wnt-signaling pathway and response to retinoic acid were observed to be activated. Collectively, this continued activity of processes involved in organ- and cellular-level development indicates ongoing developmental activity and further emphasizes the reports of developmental delay in ADHD^6, 103, 104^ and new research indicating subcortical maturation delay^105^. Secondly, the activation of ECM organization and cell-cell adhesion processes reflects ongoing or altered tissue architectural processes and cellular connectivity. These can further influence neuronal connectivity, synaptic organization, glial interactions and BBB structures and associated processes. Thirdly, immune and inflammatory responses were activated, including activation and differentiation of immune cells.

Fourthly and surprisingly, we observed overrepresentation and activation of processes associated with both ciliary structural assembly and motility. Although these findings may initially appear unexpected when considered solely in the context of the caudate, but considering the anatomical relationship between the caudate nucleus and the lateral ventricle provides a potential biological context. The caudate nucleus is not merely adjacent to the lateral ventricle; its head and body contribute directly to (more like, it forms) the lateral wall of the frontal horn and ventricular body, and its tail courses along the temporal horn^104, 106, 107^. The walls of the lateral ventricles are lined by ependymal cells, which are typically ciliated, cuboidal-to-columnar glial cells. Their cilia help move cerebrospinal fluid, while microvilli on their apical surface contribute to exchange with the CSF. This intimate anatomical relationship raises the possibility that the observed activation of biological processes related to ciliary assembly and its motility and CSF circulation from transcriptional signatures may, at least in part, suggest abnormalities in lateral ventricular functions.

Ventricular abnormalities have been reported in ADHD, but this remains a sparsely investigated area. Structural MRI studies have described alterations in lateral ventricular morphology and asymmetry^108, 109^, as well as associations between ventricular volume and neuropsychological impairment^110^. Despite these observations, the ventricular wall itself, particularly its ependymal and ciliary compartment, has received very limited attention in ADHD research. Advanced MRI approaches may improve visualization of the ventricular lining and surrounding tissue, but direct assessment of ependymal cell abundance, cellular composition, or ciliary organization generally requires tissue-based methods such as histology or microscopy. However, historical focus of ADHD research has been on neuroimaging on cortical, fronto-striatal, cerebellar, and distributed brain networks that are more directly associated with attention, executive function, motor regulation, and other behavioral manifestations of the disorder. (As ventricles don’t partake in any functions that translate into behavioral or clear physical exhibits, rather their effects are more physiological; they have not been under the limelight for exploration in ADHD.) As a result, regions with less obvious behavioral correlates, such as the ventricular system, have received comparatively less emphasis, even though their physiological roles in CSF circulation, tissue homeostasis, and brain development may still be relevant to disease biology. Large imaging studies have also increasingly shifted toward circuitry- and network-level explanations of ADHD, further contributing to the relative underrepresentation of ventricular-focused investigations. Thus, ventricular dysregulation remains largely unexplored in ADHD, and our findings highlight it as an important and previously underappreciated avenue for understanding ADHD pathophysiology. This hypothesis is further supported by prenatal studies suggesting that ventricular abnormalities can arise during fetal development and persist into childhood, potentially influencing later neurodevelopmental outcomes^111^.

By conducting MEGENA, a gene co-expression analysis, several networks of genes i.e., modules were identified that are associated with the ADHD phenotype in the caudate as well as in the putamen. Top 2,500 altered genes, including those not individually reaching statistical significance were incorporated; this broadened our lens beyond DEG-focused analyses, providing a more global view of dysregulation associated with ADHD in these brain regions. This returned all biological processes and pathways also identified by GSEA, including ciliary assembly and motility, ECM organization, and ATP metabolism and others in caudate, while additionally highlighting functions such as lipid and membrane metabolism, biosynthesis, proteostasis, and cell-cycle regulation that might have been overlooked when focusing only on statistically significant DEGs. Similarly, all biological processes identified by GSEA in the putamen, including protein stress, gliogenesis, neural remodeling, and others were also identified by MEGENA, in addition to other functions such as cytoskeletal dynamics, cell adhesion, BBB, apoptosis, and others being revealed.

In parallel with aberrant levels of metabolite regulating neurotransmitter levels (Cit), dopamine itself (i.e., neurotransmitter availability) and alterations in biological processes related to synaptic organization in caudate, its MEGENA also revealed several neuronal functions. 5 modules dominantly govern neurotransmitter production, transport to synaptic availaibility aiding neurotransmission, 2 govern neuronal connectinvity and circuit formation, 1 regulate ion homeostasis and hence membrane excitability and 1 aid in cell adhesion, ECM and BBB integrity, in addition to their other functions. In all 6 modules were identified that dominantly governed one or more neuronal functions were identified. Of these, module c1_256 emerged as the most neuronally focused gene network, with all three of its top functions related to neuronal processes including neuroglial differentiation. It consists of 10 genes, and FGD1 being the Hub regulatory gene of this network. Its parent module and sibling leaf module are also associated with ADHD, with its parent module governing 2 neuronal functions amongst its top 3 functions. This module can be considered as a candidate module for investigating neuronal alterations in the caudate and its family as a candidate family of modules.

Putamen’s MEGENA also highlighted 12 gene networks dominantly regulating neuronal functions. 3 modules each were associated with neurotransmitter production to availability at synapse and neurotransmission, membrane excitability and BBB integrity, cell adhesion and ECM; and neuroglial differentiation, neuronal connectivity and circuitry and glial functions were dominantly governed by 1 module each. Some hub genes, such as KCNE, which modulates ion flux and regulates neuronal excitability, could serve as candidate genes, and their module as candidate modules,for further exploration. These results are complementary to the biological processes revealed from GSEA analysis related to neurotransmitter reuptake i.e. recycling, GABA receptor subunit, synaptic organization as well as the dysregulated levels of dopamine.

Additionally, dysregulated levels of metabolites such as Cit and its potential relevance to glutamatergic and GABAergic neurotransmission^66, 68, 69^, reduced dopamine levels, along with the MEGENA’s identification of modules dominantly governing neuronal functions such synaptic neurotransmission, membrane e excitability and neuronal circuitry in caudate and putamen, provide strong evidence implicating dysregulation in of neurotransmitter availability at synapse in both these brain regions.

Putamen’s module c1_307 would be a candidate gene network for investigating aberrations in developmental processes, as it governs neuronal and glial differentiation and development as well as brain morphogenesis, and so does its parent module; however, as a leaf module, c1_307 is functionally more specialized in regulating these. Putamen has more modules governing brain morphogenesis and neuroglial differentiation and development than caudate, where 5 modules regulate to the prior and 8 regulate the latter. Several modules also govern morphogenesis and differentiation in the putamen as dominant functions. Additionally, differentiation, development, and morphogenesis are dominantly governed processes of most of caudate’s modules (10 out of 25 modules). Not as dominant function but neuroglial differentiation was observed as additional functions of 6 caudate modules. Both these regions exhibit strong signs of altered developmental processes. These suggest developmental processes are prominently affected in both these brain regions. These findings are consistent with previous neuroimaging evidence of delayed cortical maturation in children and lower brain volume with ADHD^6, 7^.

Most of putamen’s modules regulated immune and inflammatory processes (19 out of 33 modules). Several modules also dominantly governed membrane metabolism, signaling cascades, cell cycle and surveillance and metabolism/ biosynthesis and differentiation and development related functions. In addition to these, few modules were also dominantly associated to proteostasis (including synthesis, QC and degradation), intracellular trnasport, DNA damage repair and endocrine signaling. On the other hand, Most of the modules identified in the caudate are associated with ADHD-regulated differentiation, development (10 out of 25 modules) and morphogenesis in their top 3 functions, indicating signs of continued or altered development through later stages of adolescence. Several modules also dominantly regulated signaling cascades, immune and inflammatory responses, and lipid/ membrane metabolism. Some modules were also linked to, cellular homeostasis, cell cycle and surveillance, energy metabolism, and ciliary motility, in addition to other functions that were dominantly governed by lesser modules. Most functions identified for modules, are in line with the processes observed in enrichment analysis. For instance, the modules related to intracellular trafficking in both regions also link to the shared down-regulated genes’ enrichment for ER-Golgi vesicle transport. The gene networks and their regulatory hub genes provide not only evidence but also an avenue to explore these less-linked biological functions in ADHD.

A final key aspect of this study was the cross-study comparison, which assessed the consistency of transcriptomic alterations across independent datasets. Comparison of our caudate DEGs with the only available bulk RNA-seq caudate study by Sudre et al.^43^ identified only 35 overlapping DEGs, most with opposing effect directions, indicating limited concordance at the individual-gene level. Differences in cohort design, especially the difference in age was potentially the major contributing factor to this variability, as our study focused on early developmental age range till 19 years with clinically diagnosed combined or inattentive ADHD, whereas the other study included a broader age range (6.7–38 years) without subtype restriction; additional clinical, biological, and technical factors may also contribute. This variability in findings owing to age difference is line with abberations observed in brain volumes at young age which disappear in adults especially in caudate^19, 20^ and supports our hypothesis that underlying alterations in ADHD vary with age. Meta-analysis identified 809 significant DEGs, of which 409 showed concordant effect directions and heterogeneity ≤5%, representing more consistent cross-study signals, irrespective of age and subtype differences in study designs. Under the most stringent criterion, five genes—AQP7, CALHM1, PRRC2A, NPIPB5, and TTC23-AS1—were significant in both individual studies and the meta-analysis representing most conservative set of genes. Together, these findings highlight substantial gene-level variability across studies while demonstrating the value of meta-analysis and suggesting that future ADHD transcriptomic studies may benefit from greater developmental and clinical homogeneity or stratification by age and subtype.

Furthermore, we also examined the expression of 76 ADHD risk genes reported by Demontis et al.^12^ across the caudate and putamen. Only ARTN, a glial cell line-derived neurotrophic factor involved in dopaminergic neuronal survival, axon guidance, differentiation regeneration and neuridevelopment^112, 113, 114^, was significantly altered, showing a 3.46-fold upregulation in the putamen, while none of the 76 genes were significantly dysregulated in the caudate. This limited overlap between GWAS-prioritized genes and brain expression changes further underscores the need for transcriptomic, metabolomic, and other molecular studies to bridge the gap between genetic risk and molecular dysregulation in the ADHD brain.

To summarize, we identify 3 broader functional alterations found in both of these brain regions. Firstly, several metabolites, genes, biological processes, and module functions have consistently provided considerable evidence highlighting alterations in neurotransmitter availability and neurotransmission. Additionally, perturbations found in energy (ATP) metabolism/ synthesis also directly could affect neurotransmission. Secondly, immune and inflammatory responses were another dysregulated functions, and their alterations potentially be linked to reduced levels of dopamine as well as ventricular perturbations. This function had more prominent dysregulation in putamen. Developmental processes related to differentiation and morphogenesis revealed from module functions as well as their up-regulation in both brain regions in enrichment analysis implicate developmental delay which has been strongly associated to ADHD. This study suggests a prolonged developmental delay beyond childhood through adolescence phase. However, these do not indicate convergent dysregulation, as some processes showed alterations in opposite directions and only 2.64% of gene-level overlap highlight strong region-specific alterations even within anatomically related components of the striatum. In addition, thispaper also reveals several other cellular processes that potentially could be linked to ADHD. It also provides several gene networks and dysregulated genes for more focused explorations of pathways and functions at cellular and molecular levels for future investigation. Additionally, it provides a large subset of genes potentially consistently deregulated across childhood to adulthood developmental stages and across ADHD subtypes. This paper highlights and opens several avenues of investigating cellular amd molecular basis underlying ADHD. Furthermore the paper, highlights the need for reducing heterogeneity in sample cohorts, especially in age and subtype aspects of ADHD to reduce variability in findings and emphasizes the need for expression level studies to bridge the exisiting gap in our understanding of this disorder.

## Code Availability

All code scripts used for data processing, statistical analysis, and visualization will be made available upon request by the corresponding or the first author.

## Supporting information

All Supplementary Figures

All Supplementary Tables

Additional information-caudate's modules

Additional information-putamen's modules

## Acknowledgements

The tissue used in this research was obtained from the Human Brain Collection Core, Intramural Research Program, NIMH (http://www.nimh.nih.gov/hbcc). Gemini api was used for sorting of GO terms (from enrichment results of MEGENA modules) into 29 functional themes. ChatGPT was used to assist with grammar and phrasing of sentences.

