## Supplementary material for "Transcriptomic and Metabolomic Analysis of the Caudate and Putamen in ADHD": All Supplementary Figures: Rajvi_ADHD_Supplementary_Figures.pdf

### Supplementary Figure 1:

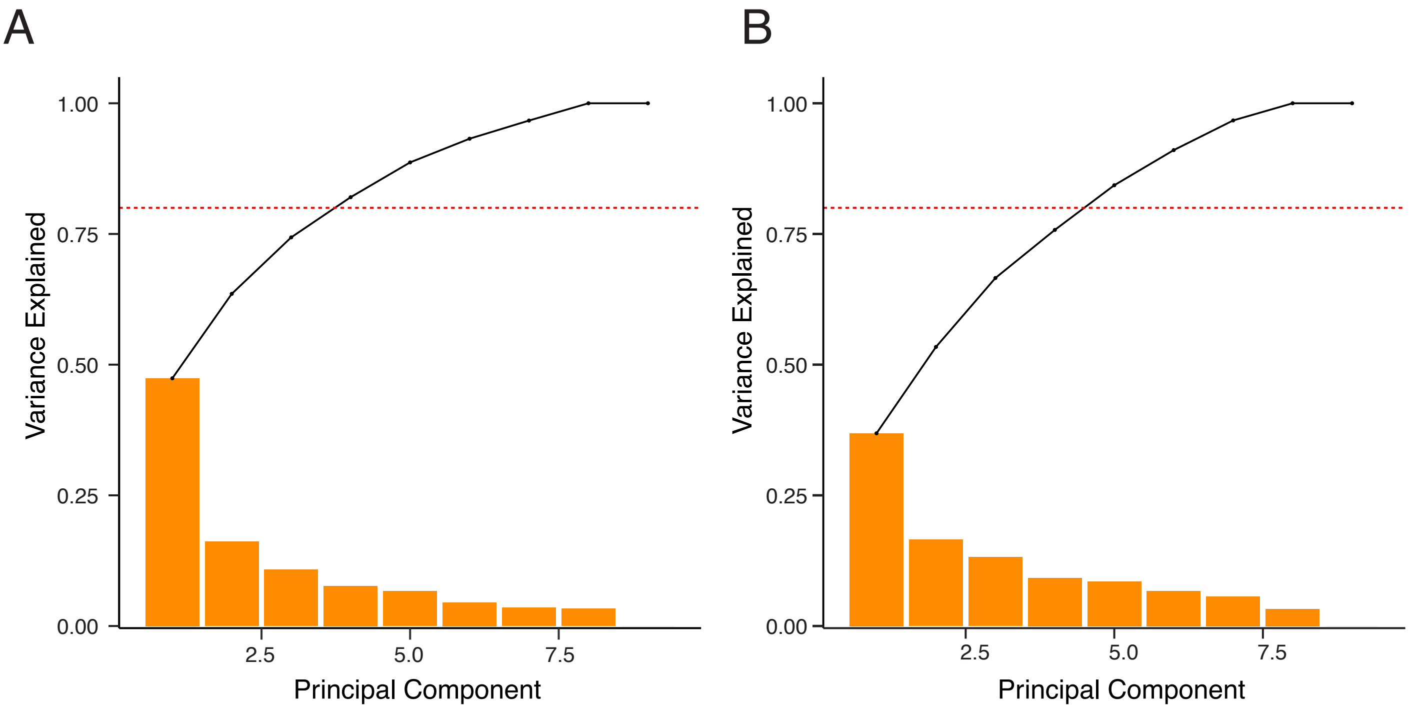


Supplementary Figure 1: PCA plots of Bulk RNA-sequencing samples: (A, B) PCA bar graphs showing the percentage of variance explained by each PC, the dashed line indicates first 4 PCs reaching approximately 80% as a reference.

### Supplementary Figure 2:

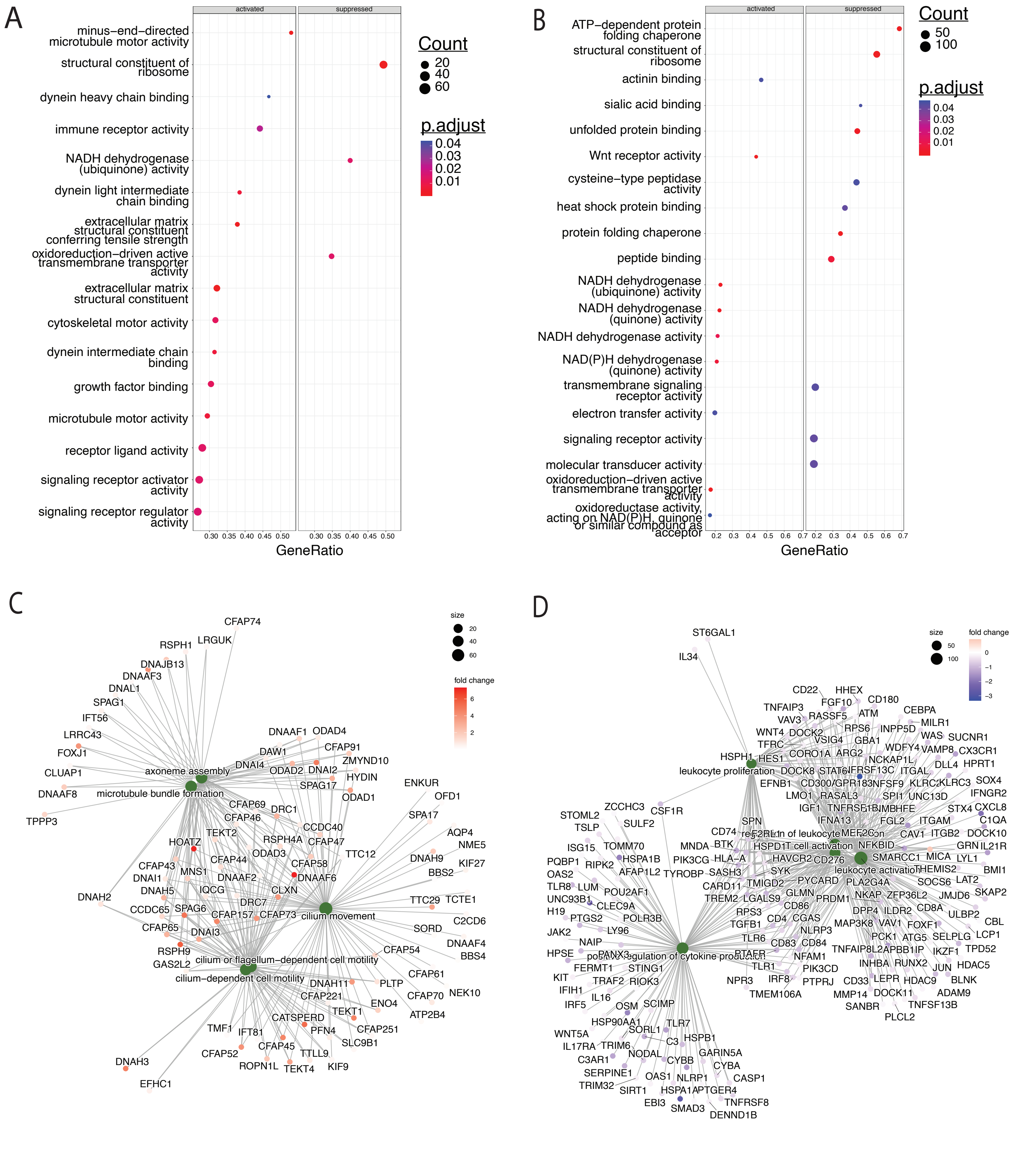


Supplementary Figure 2: Functional consequence of Caudate and Putamen's DEGs (A, B) Molecular functions affected by significant DEGs in our Caudate (A) and Putamen (B) datasets respectively and (C, D) Network plots representing top 5 BPs and their associated significant DEGs as analyzed from our Caudate (C) and Putamen (D) datasets respectively.

### Supplementary Figure 3:

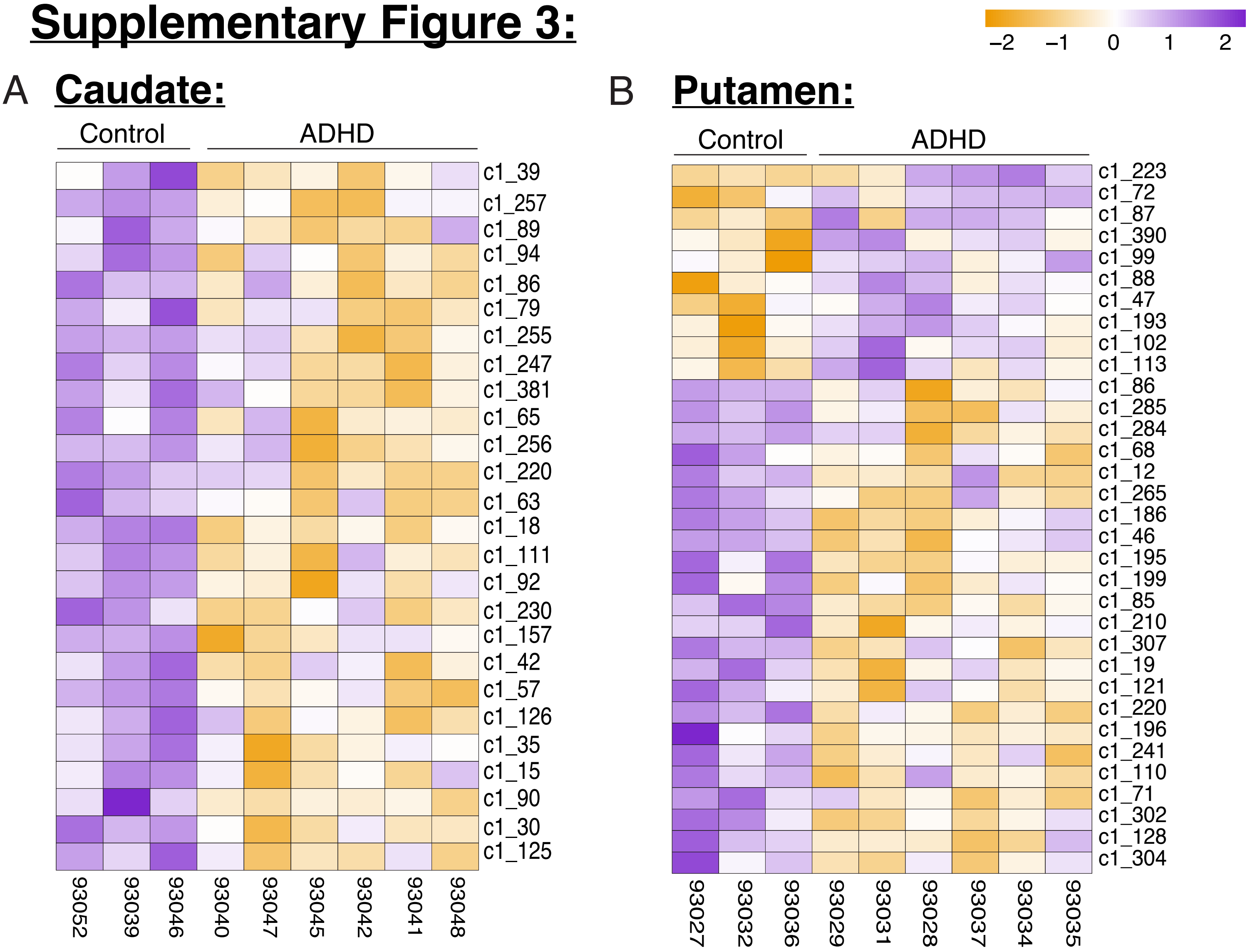


Figure 3: Module eigengene scores of modules from MEGENA analysis: (A, B) Module eigengene (ME) scores of each module (list as rows in the matrix) from every sample (list as columns in the matrix) formed of genes created based on co-expression pattern of top 2500 DEGs in caudate (A) and putamen (B) tissues respectively. The samples listed as columns are grouped where first 3 belong to control cohort and the rest 6 belong to ADHD cohort.

### Supplementary Figure 4:

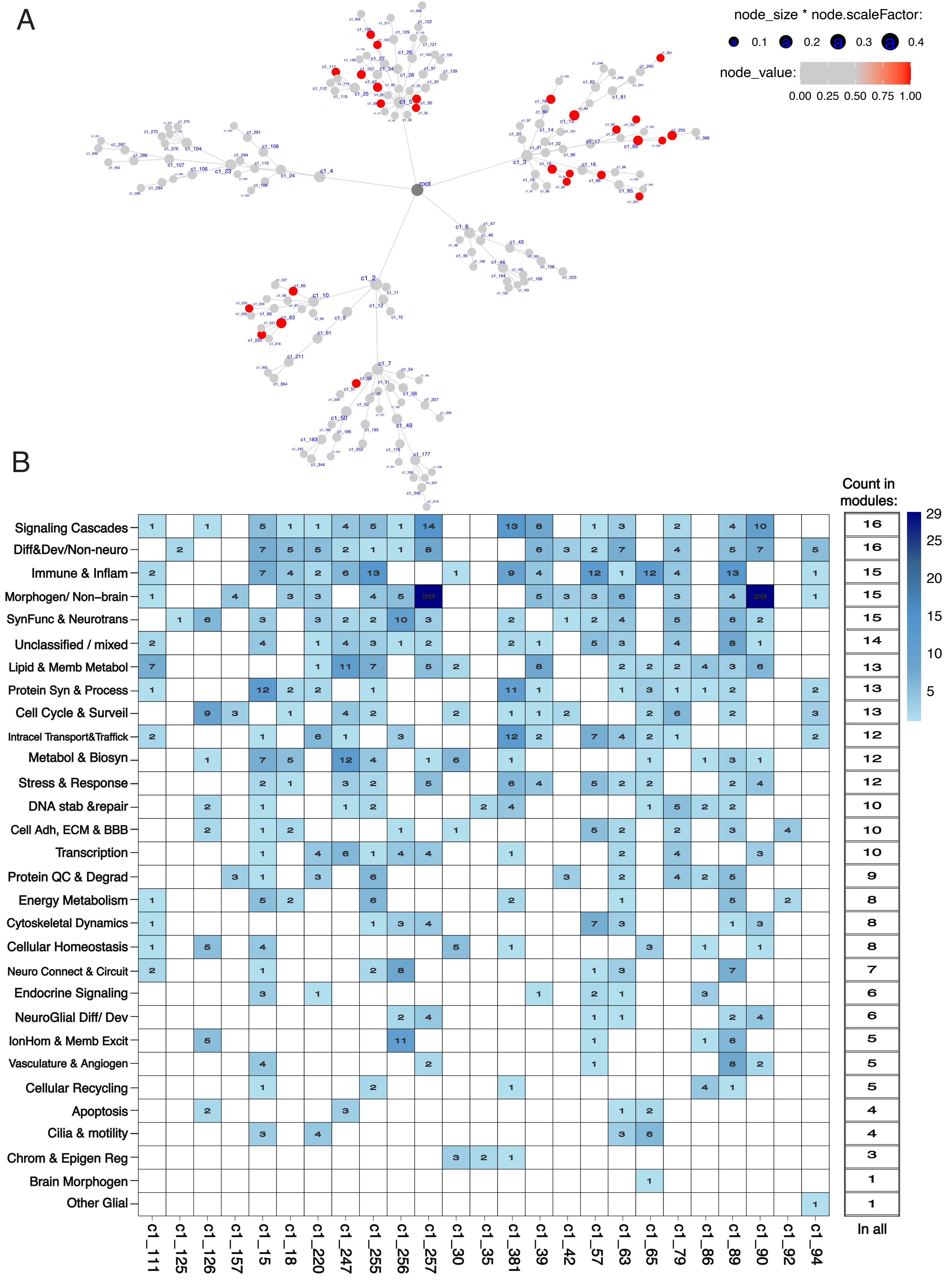

Figure 4: Hierarchy and functions of Modules of Caudate tissue: (A) Hierarchal plot of all the modules created from MEGENA analysis of top 2500 DEGs from caudate tissue showing the connection between the modules, with every branching the genes get divided and grouped with genes that they are more closely related to in terms of their co-expression patterns. The modules in red have significant assocaition with ADHD as phenotype and the size of circle is directly proportional to the module size i.e. the number of genes thatmake up the module. (B) Heatmaps representing all the modules of caudate and all the functional biological themes assocaited with each module's genesets, with the heatmap tiles showing the number of GO terms assigned to that functional theme for that module. The biological themes are represented in acronyms with the guide to their full names mentioned in Appendix/ Table #. The column on the right of the heatmap mentions for the number of modules that have that biological theme as its associated function.

### Supplementary Figure 5:

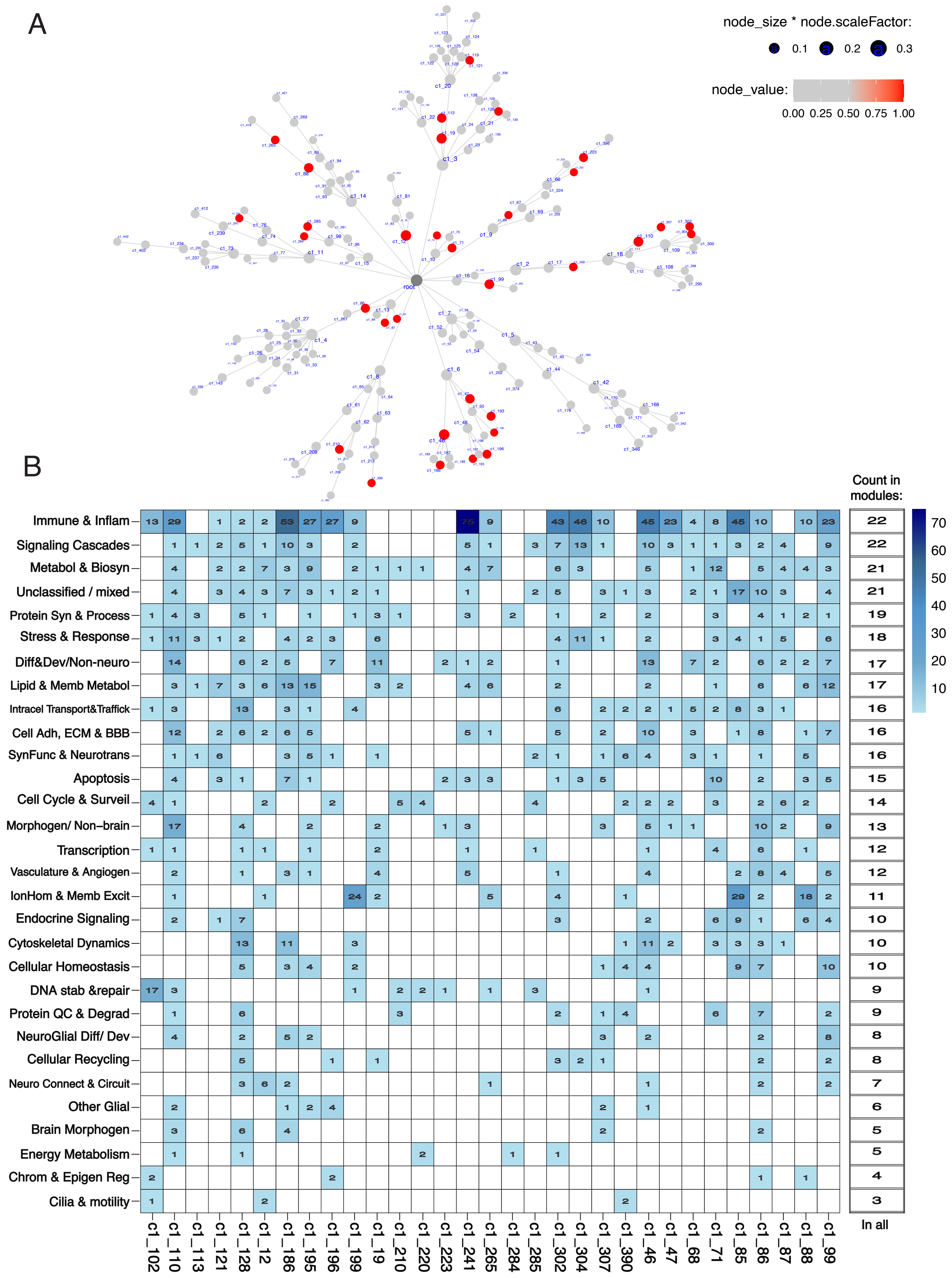

Figure 5: Hierarchy and functions of Modules of Putamen tissue: (A) Hierarchal plot of all the modules created from MEGENA analysis of top 2500 DEGs from putamen tissue showing the connection between the modules, with every branching the genes get divided and grouped with genes that they are more closely related to in terms of their co-expression patterns. The modules in red have significant assocaition with ADHD as phenotype and the size of circle is directly proportional to the module size i.e. the number of genes that make up the module. (B) Heatmaps representing all the modules of putamen and all the functional biological themes assocaited with each module's genesets, with the heatmap tiles showing the number of GO terms assigned to that functional theme for that module. The biological themes are represented in acronyms with the guide to their full names mentioned in Appendix/ Table #. The column on the right of the heatmap mentions for the number of modules that have that biological theme as its associated function.

#### Supplementary Figure 6:

##### Caudate_c1_89

A

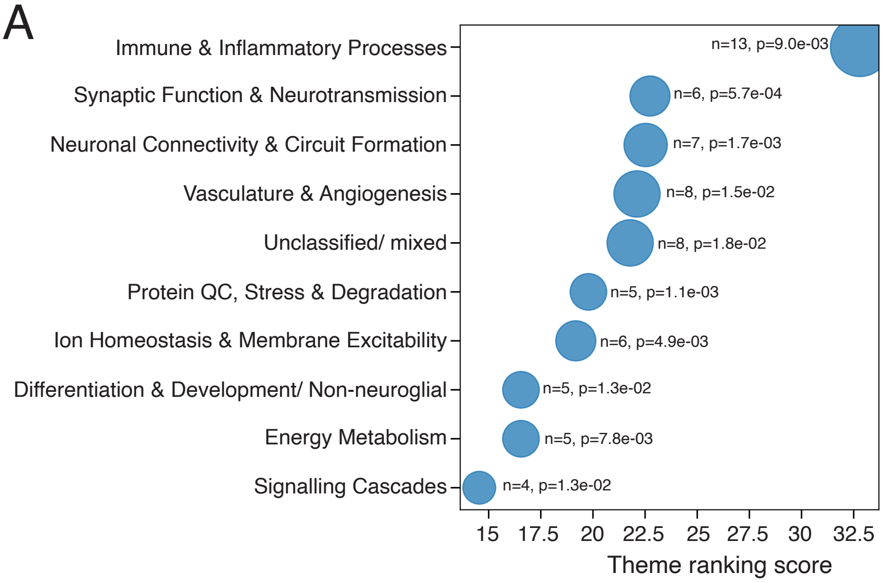


B

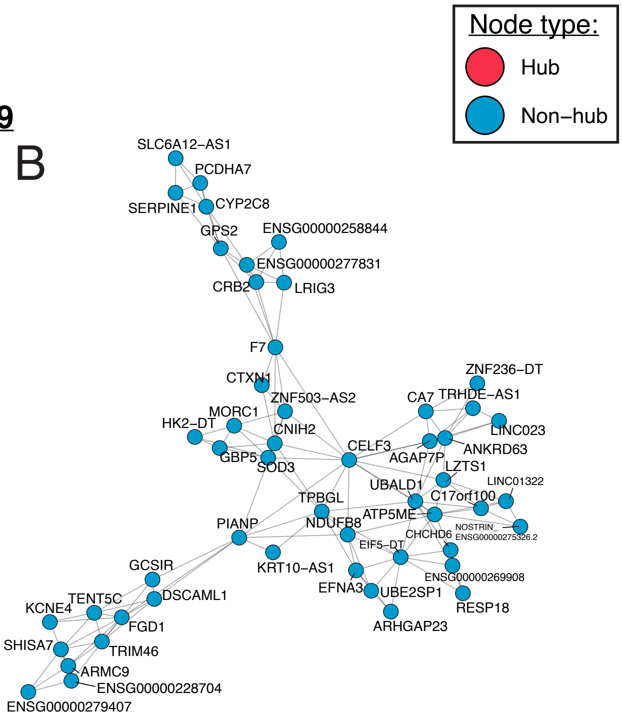


##### Caudate_c1_255

C

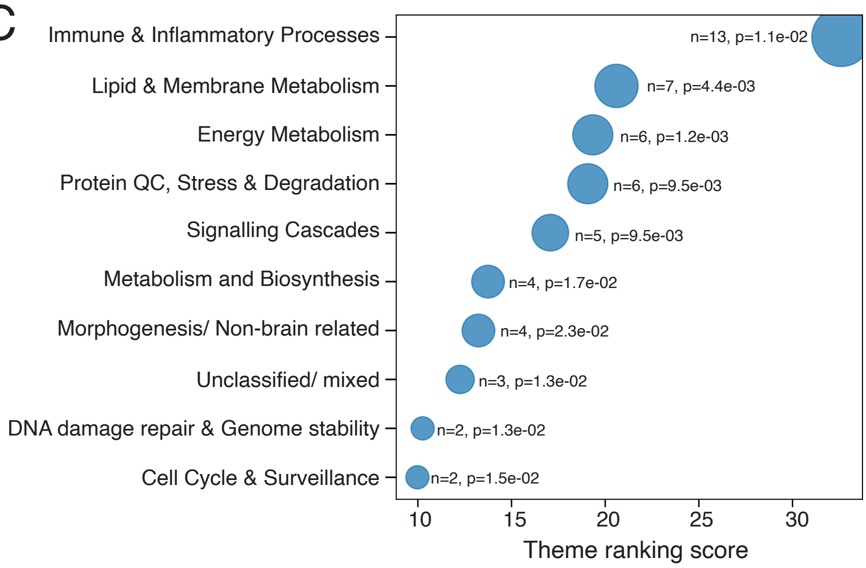


D

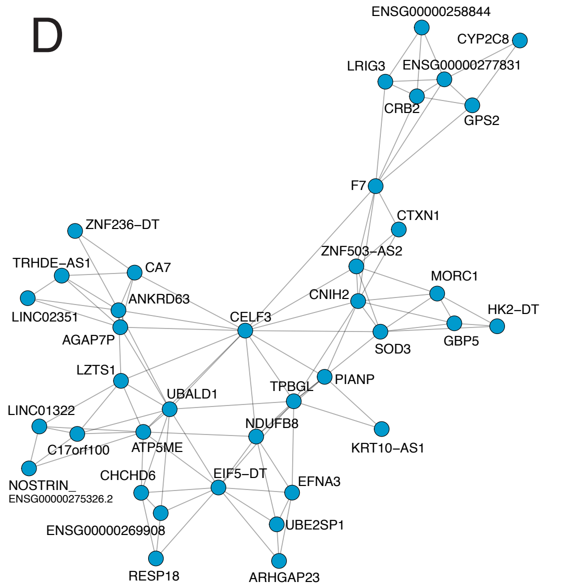


E

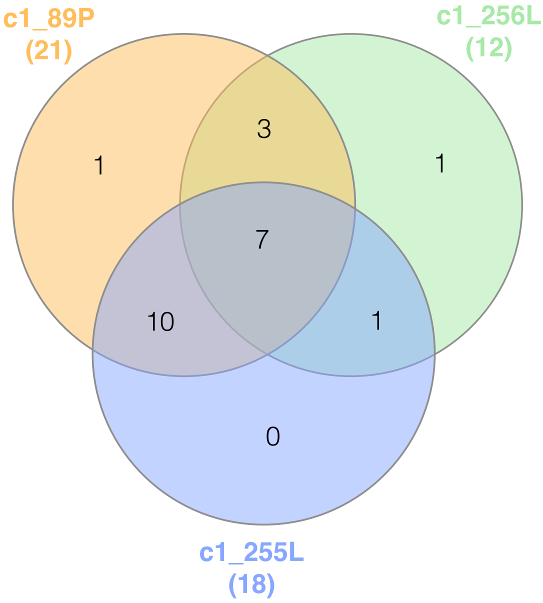


F

| All (c1_89P, c1_256L & c1_255L) | c1_89P & c1_255L | c1_89P & c1_256L | c1_256L & c1_255L |
|---|---|---|---|
| SynFunc & Neurotrans | Immune & Inflam | IonHom & Memb Excit | Transcription |
| Neuro Connect & Circuit | Protein QC & Degrad | Cell Adh, ECM & BBB | |
| Unclassified/ mixed | Energy Metabolism | NeuroGlial Diff/ Dev | |
| Diff&Dev/Non-neuro | Protein Syn & Process | | |
| Signaling Cascades | Lipid & Memb Metabol | | |
| Morphogen/ Non-brain | Metabol & Biosyn | | |
| Cytoskeletal Dynamics | DNA stab & repair | | |
| | Cell Cycle & Surveil | | |
| | Stress & Response | | |
| | Cellular Recycling | | |

Supplementary Figure 6: Functional characterization of parent-leaf modules significanlty associated with ADHD in Caudate: (A, C) The functional association of parent module c1_89 (A) and its leaf module c1_255 (C) modules' genesets through representation of their top 10 biological themes in Dot plot. Each biologocal theme classifies the very specific and precise Gene Ontology (GO) terms of Biologiocal Processes (BPs) from EnrichR results, into broader and interpretable terms. The themes are ranked by a ranking metric of: Rank = 3(-log10p-value) + 2n + combined score, where n is the number of GO terms in the theme, -log10p-value of the top BP in the theme and its combined score both as calculated by EnrichR. y-axis indicates the theme names, the ranking score for each theme is plotted on x-axis, n and p-value are mentioned besides the dot for each theme and the size of ech dot is relative to its n. (B, D) Network plots of modules c1_89 (B) and c1_255 (D) with the genes constituting the module and the connections between those genes. (E) Venn diagram representing the the number of shared and unique functional themes between parent module c1_89 and its leaf modules c1_255 and c1_256 (represented in Figure 2) and (F) table showing the lists of shared functional biological themes.

#### Supplementary Figure 7:

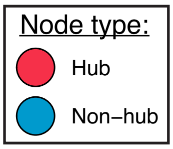


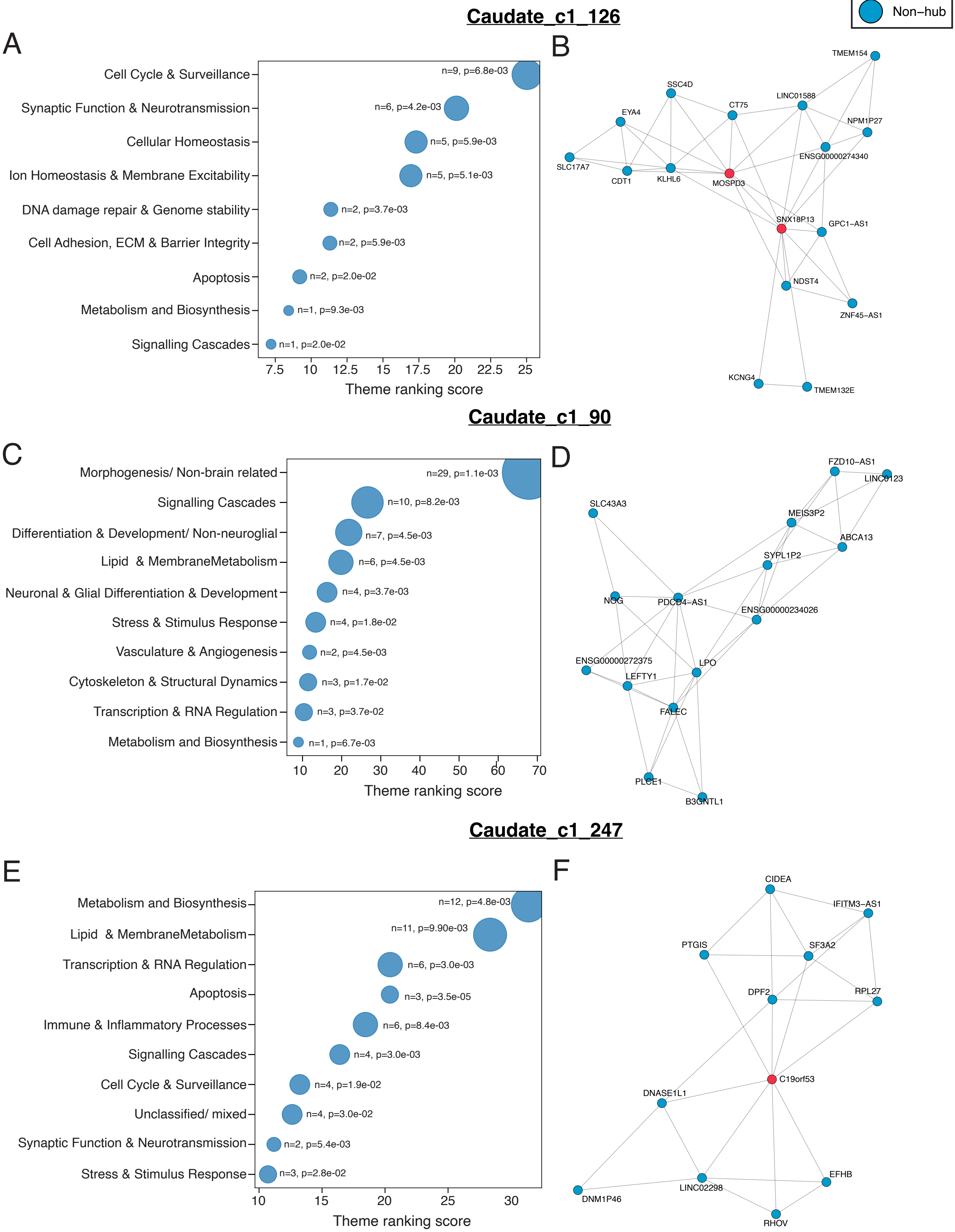

Supplementary Figure 7: Functional characterization of modules with different roles and modue sizes but significanlty associated with ADHD in Caudate: (A, C, E) The functional association of modules c1_126 (A), c1_90 (C) and c1_247 (E) modules' genesets through representation of their top 10 biological themes in Dot plot. y-axis indicates the theme names, the ranking score for each theme is plotted on x-axis, n and p-value are mentioned besides the dot for each theme and the size of ech dot is relative to its n. (B, D, F) Network plots of modules c1_126 (B), c1_90 (D) and c1_247 (F) with the genes constituting the module and the connections between those genes and the Hub gene (highlighted in red) which are the highly connected genes within the module.

### Supplementary Figure 8:

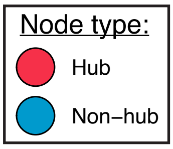


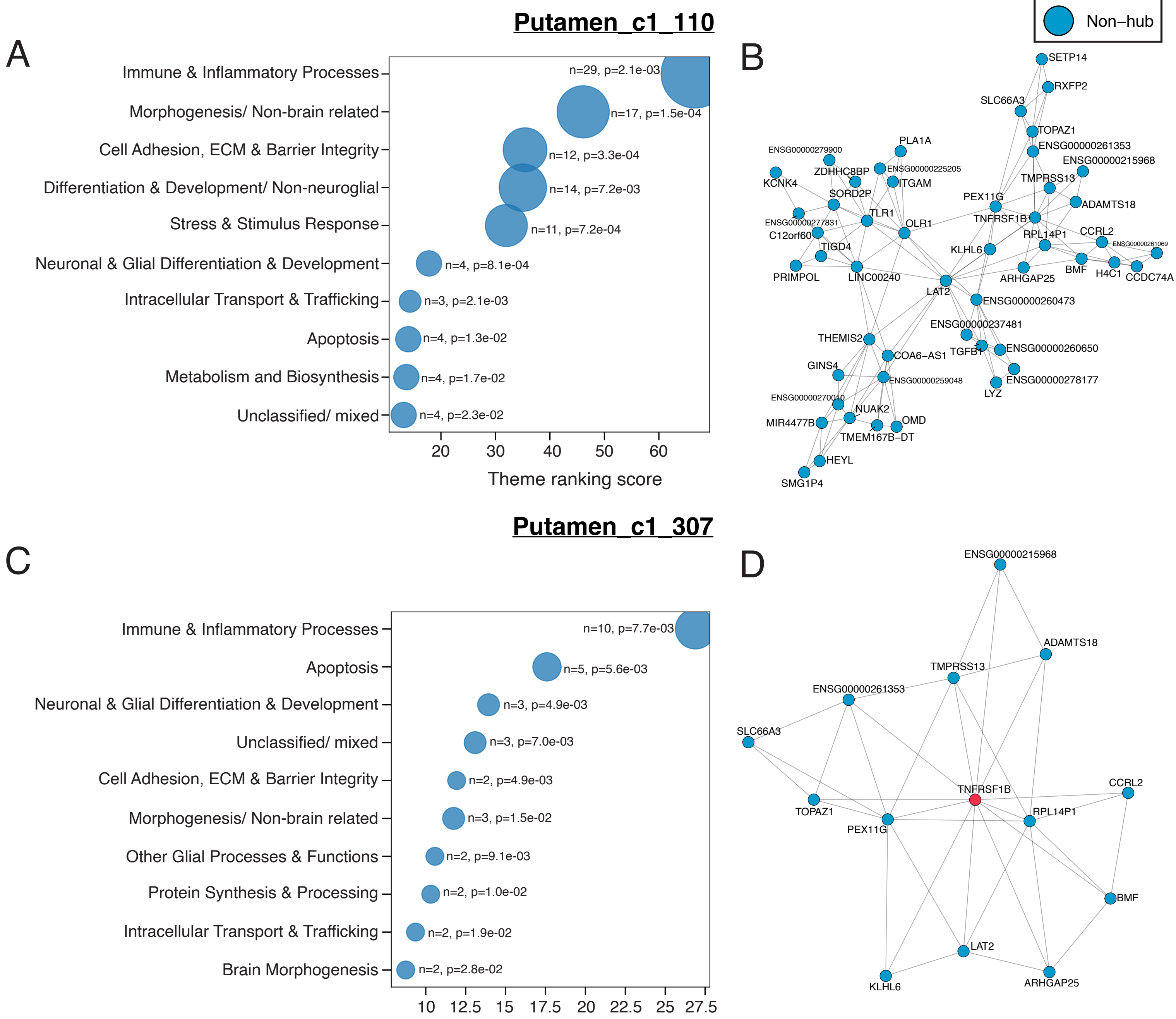


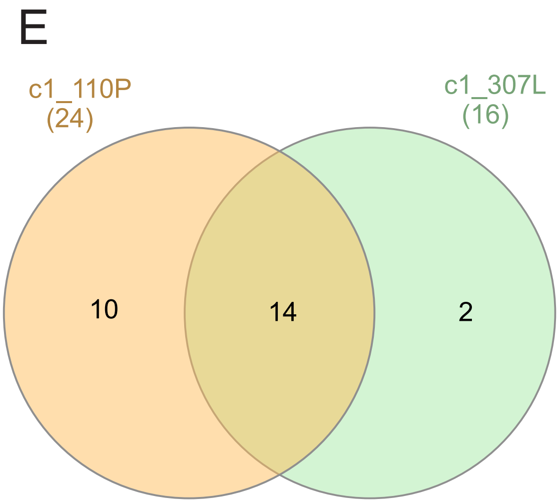


F

| c1_110P | c1_110P & c1_307L | c1_307L |
|---|---|---|
| Immune & Inflammatory Processes | Immune & Inflammatory Processes | Immune & Inflammatory Processes |
| Morphogenesis/ Non-brain related | Morphogenesis/ Non-brain related | Apoptosis |
| Cell Adhesion, ECM & Barrier Integrity | Cell Adhesion, ECM & Barrier Integrity | Neuronal & Glial Differentiation & Development |
| Differentiation & Development/ Non-neuroglial | Stress & Stimulus Response | Unclassified / mixed |
| Stress & Stimulus Response | Neuronal & Glial Differentiation & Develc | Cell Adhesion, ECM & Barrier Integrity |
| Neuronal & Glial Differentiation & Development | Intracellular Transport & Trafficking | Morphogenesis/ Non-brain related |
| Intracellular Transport & Trafficking | Apoptosis | Other Glial Processes and Functions |
| Apoptosis | Unclassified / mixed | Protein Synthesis & Processing |
| Metabolism and Biosynthesis | Brain Morphogenesis | Intracellular Transport & Trafficking |
| Unclassified / mixed | Protein Synthesis & Processing | Brain Morphogenesis |
| Brain Morphogenesis | Other Glial Processes and Functions | Synaptic Function & Neurotransmission |
| Protein Synthesis & Processing | Synaptic Function & Neurotransmission | Cellular Homeostasis |
| Lipid & Membrane Metabolism | Protein QC, Stress & Degradation | Signaling Cascades |
| DNA damage repair & Genome stability | Signaling Cascades | Protein QC, Stress & Degradation |
| Vasculature & Angiogenesis | | Stress & Stimulus Response |
| Other Glial Processes and Functions | | Cellular Recycling |
| Endocrine & Neuropeptide Signaling | | |
| Synaptic Function & Neurotransmission | | |
| Cell Cycle & Surveillance | | |
| Transcription & RNA Regulation | | |
| Energy Metabolism | | |
| Ion Homeostasis & Membrane Excitability | | |
| Protein QC, Stress & Degradation | | |
| Signaling Cascades | | |

Supplementary Figure 8: Functional characterization of parent-leaf modules significanlty associated with ADHD in Putamen: (A, C) The functional association of parent module c1_110 (A) and its leaf module c1_307 (C) modules' genesets through representation of their top 10 biological themes in Dot plot. y-axis indicates the theme names, the ranking score for each theme is plotted on x-axis, n and p-value are mentioned besides the dot for each theme and the size of ech dot is relative to its n. (B, D) Network plots of modules c1_110 (B) and c1_307 (D) with the genes constituting the module and the connections between those genes and the Hub gene (highlighted in red) which are the highly connected genes within the module. (E) Venn diagram representing the the number of shared and unique functional themes between parent module c1_110 and its leaf modules c1_307 and (F) table showing the lists of their shared and unque functional biological themes.

#### Supplementary Figure 9:

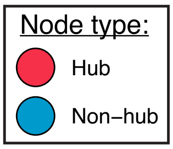


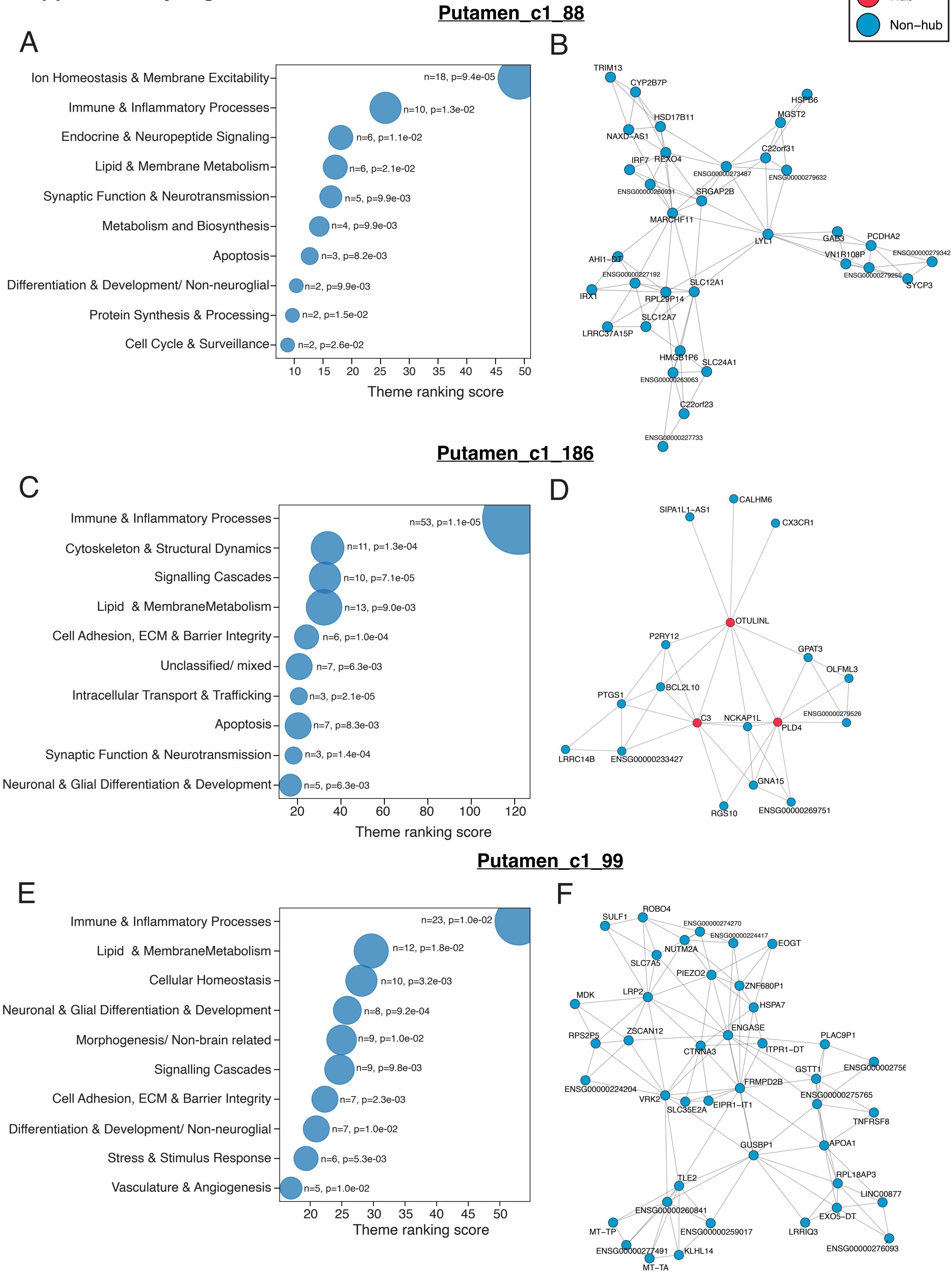

Supplementary Figure 9: Functional characterization of modules with different roles and modue sizes but significanlty associated with ADHD in Putamen: (A, C, E) The functional association of modules c1_88 (A), c1_186 (C) and c1_99 (E) modules' genesets through representation of their top 10 biological themes in Dot plot. y-axis indicates the theme names, the ranking score for each theme is plotted on x-axis, n and p-value are mentioned besides the dot for each theme and the size of ech dot is relative to its n. (B, D, F) Network plots of modules c1_88 (B), c1_186 (D) and c1_99 (F) with the genes constituting the module and the connections between those genes and the Hub gene (highlighted in red) which are the highly connected genes within the module.

#### Supplementary Figure 10:

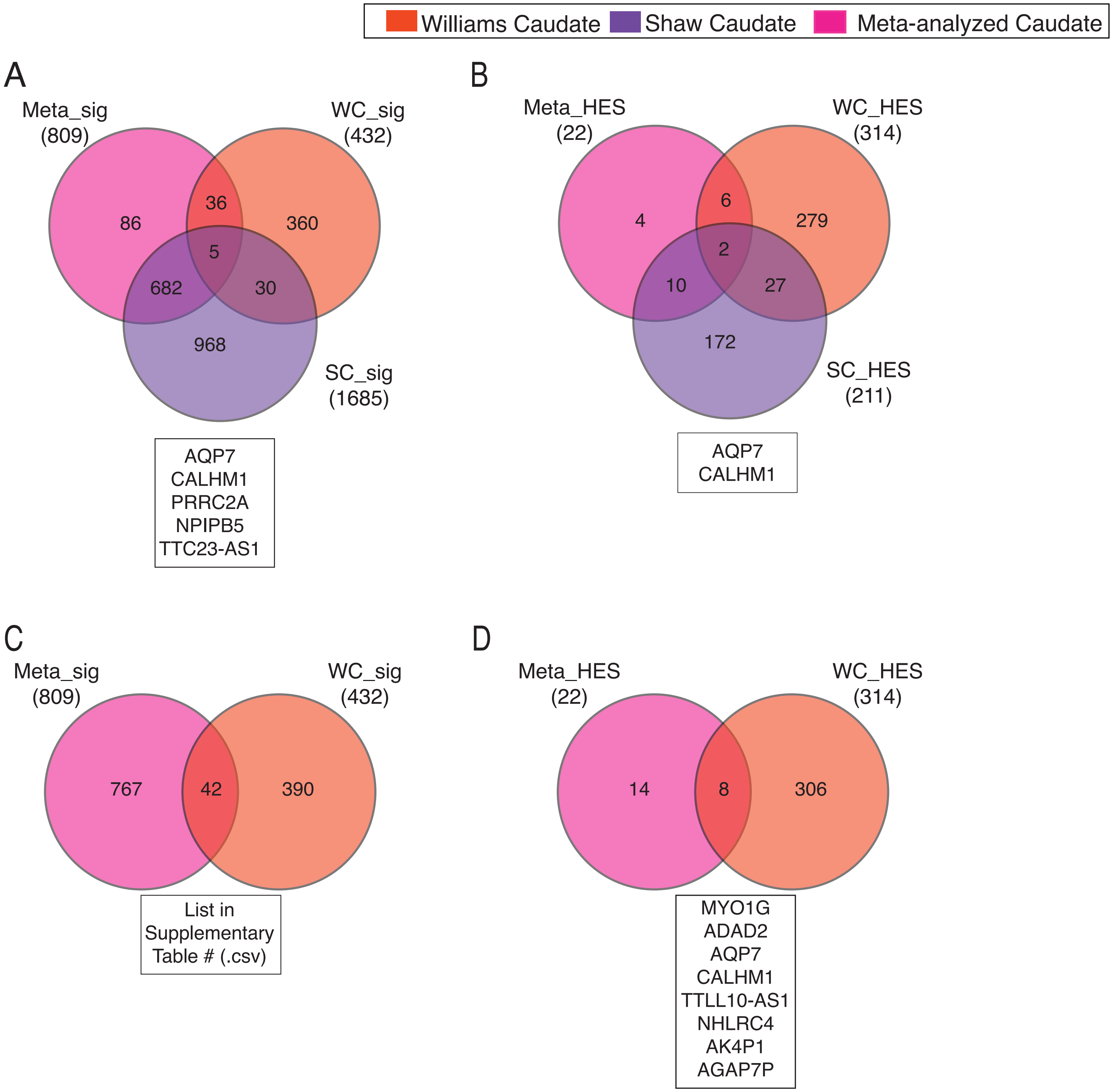


Supplementary Figure 10: Shared Differetially Expressed Genes of Caudate between studies/ analyses: (A, C)Statistically significant DEGs shared between (A) Meta-analyzed Caudate, this study's Caudate and Sudre et al. study's Caudate, all 3 datasets, and (C) Meta-analyzed and this study's Caudate datasets, (B, D) Highly Expressed significant (HES) DEGs between (B) Meta-analyzed, this study's and Sudre et al.'s Caudate, all 3 datasets, and (D) Meta-analyzed and this study's Caudate datasets. (significant DEGs are the ones with p-value =< 0.05).

#### Supplementary Figure 11:

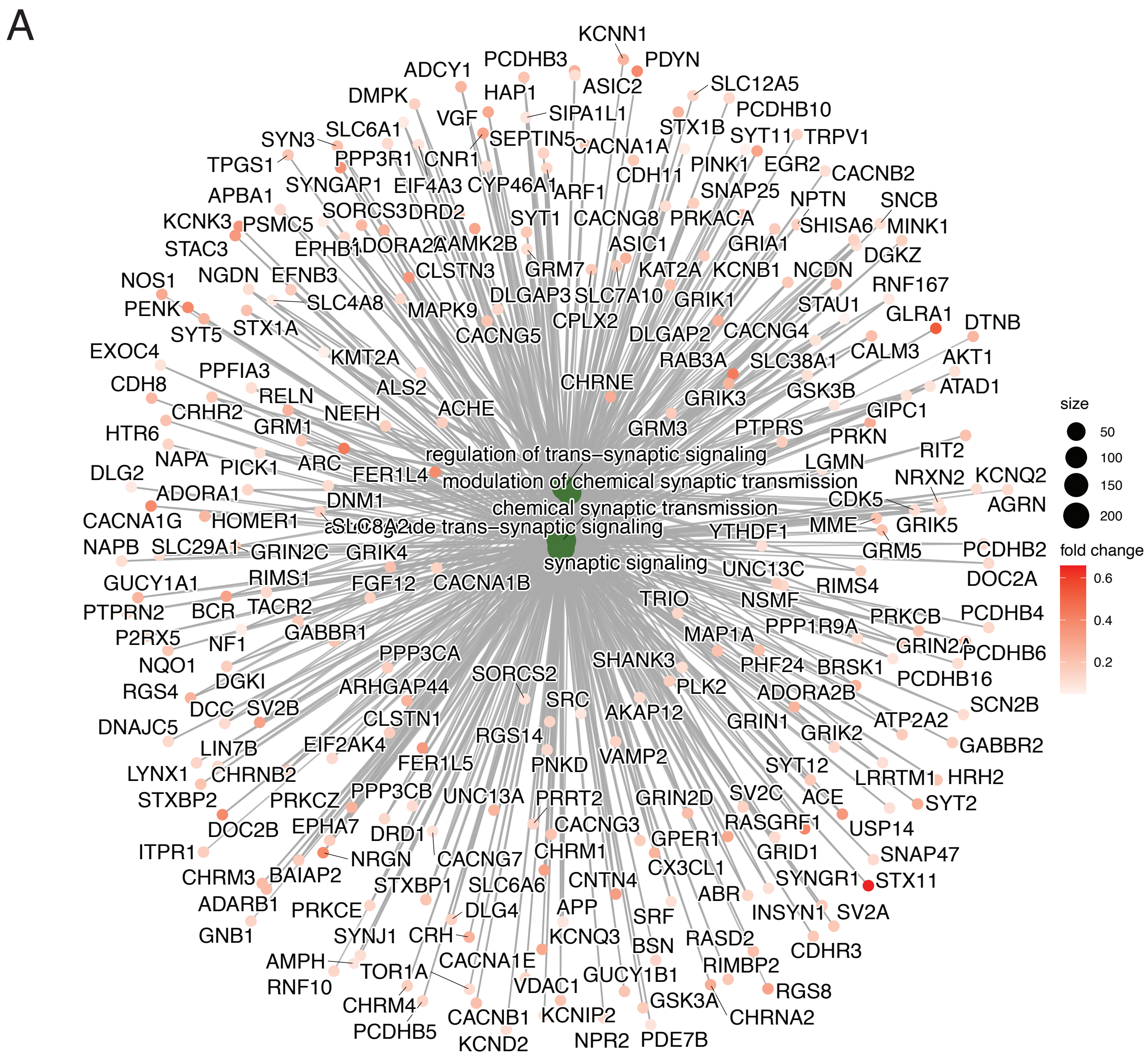


Supplementary Figure 11: Functional association of DEGs identified from Meta-analyzed caudate dataset: (A) Network plot highlighting the top 5 Biological Processes (BPs) identified from GSEA anallysis of meta-analyzed caudate dataset's DEGs and all the DEGs from that set that are associated with these top 5 BPs.

### Supplementary Figure 12:

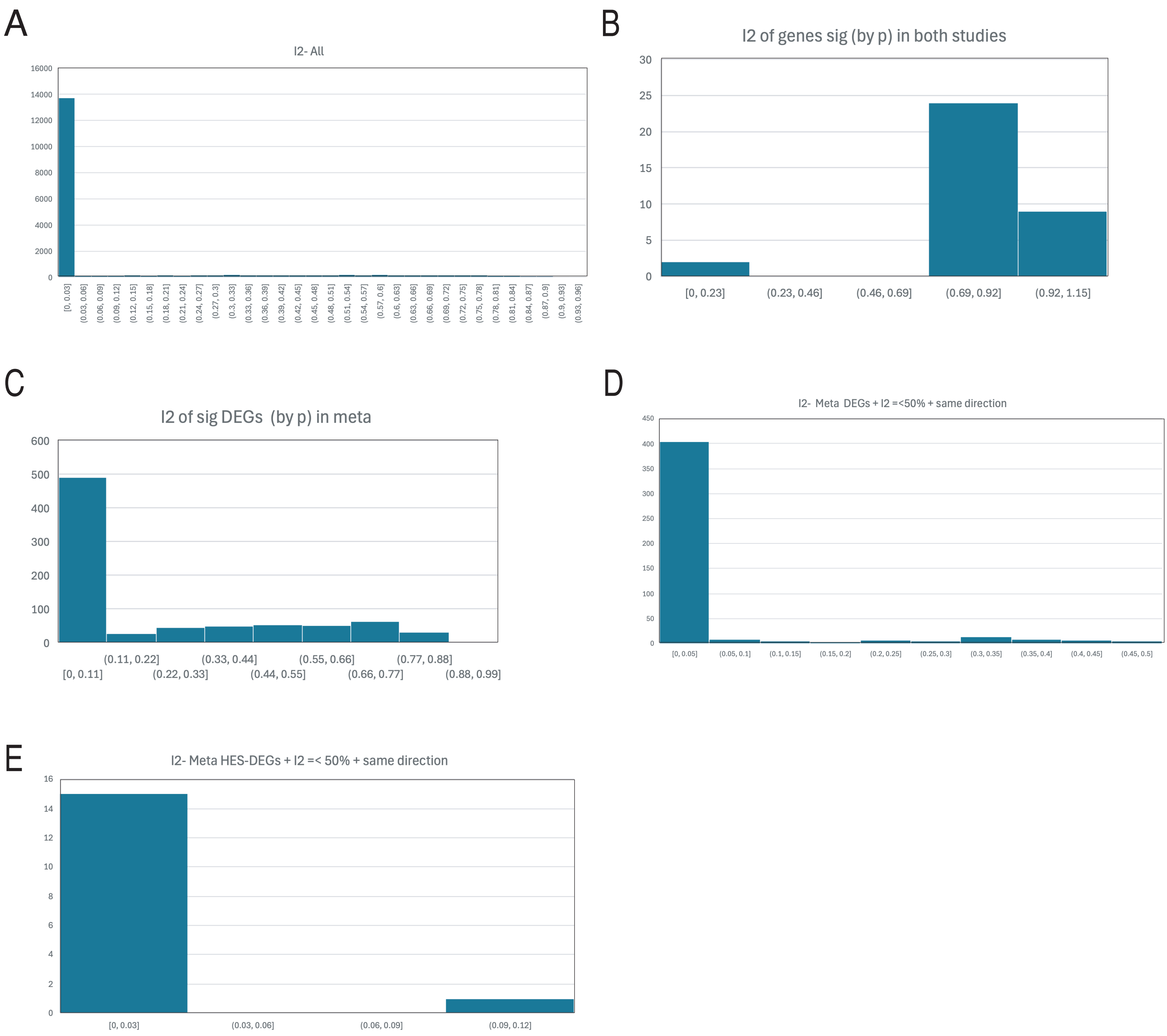


<u>Supplementary Figure 12: Analyzing $I^2$ of different gene subsets caudate's Meta-anlaysis to assess heterogeneity between 2 caudate's Bulk RNA-seq datasets:</u> Histograms showing distribution of genes across $I^2$ ranges for (A) all genes with meta-analyzed results, (B) genes overlapping between the 2 bulk RNA-sequencing datasets individually, (C) all significant DEGs identified through Meta-analysis, (D) Meta-analysis significant DEGs with $I^2$ =<0.5 (i.e. 50%) and had same direction of change in indivudal datatsets, and (E) Metaanalysis HES-DEGs with $I^2$ =<0.5 (i.e. 50%) and had same direction of change in indivudal datatsets, study's Caudate datasets. (significant DEGs are the ones with p-value =< 0.05). Here the numbner of genes is represented on y-axis, $I^2$ ranges on x-axis and bars indicat the number of genes per bin (i.e. $I^2$ range).
